# Investigating tissue microstructure using steady-state diffusion MRI

**DOI:** 10.1101/2024.05.15.594140

**Authors:** Benjamin C. Tendler

**Affiliations:** Wellcome Centre for Integrative Neuroimaging, FMRIB, Nuffield Department of Clinical Neuroscience, University of Oxford, Oxford, UK

## Abstract

Diffusion MRI is a leading method to non-invasively characterise brain tissue microstructure across multiple domains and scales. Diffusion-weighted steady-state free precession (DW-SSFP) is an established imaging sequence for post-mortem MRI, addressing the challenging imaging environment of fixed tissue with short T_2_ and low diffusivities. However, a current limitation of DW-SSFP is signal interpretation: it is not clear what diffusion ‘regime’ the sequence probes and therefore its potential to characterise tissue microstructure. Building on a model of Extended Phase Graphs (EPG), I establish two alternative representations of the DW-SSFP signal in terms of (1) conventional b-values (time-*independent* diffusion) and (2) encoding power-spectra (time-*dependent* diffusion). The proposed representations provide insights into how different parameter regimes and gradient waveforms impact the diffusion properties of DW-SSFP. Using these representations, I introduce an approach to incorporate existing diffusion models into DW-SSFP without the requirement of extensive derivations. Investigations incorporating free-diffusion and tissue-relevant microscopic restrictions (cylinder of varying radius) give excellent agreement to complementary analytical models and Monte Carlo simulations. Experimentally, the time-*independent* representation is used to derive Tensor and proof of principle NODDI estimates in a whole human post-mortem brain. A final SNR-efficiency investigation demonstrates the theoretical potential of DW-SSFP for ultra-high field microstructural imaging.

## Introduction

Establishing non-invasive neuroimaging technologies to characterise tissue microstructure facilitates investigations into the human brain and the efficacy of novel neurotherapeutic targets. Diffusion MRI is a leading neuroimaging modality in this space, probing tissue microstructure by sensitising images to changes in cellular morphology. At present, diffusion MRI investigations are almost exclusively performed using the diffusion-weighted spin echo (DW-SE) MRI sequence^[1]^ (Figure 1a). DW-SE provides an intuitive framework to understand and investigate diffusion across multiple domains and scales. More advanced diffusion acquisition schemes^[2]^ are often fundamentally based on the DW-SE, making it the principal technique to non-invasively characterise tissue microstructure.

**Figure 1:**
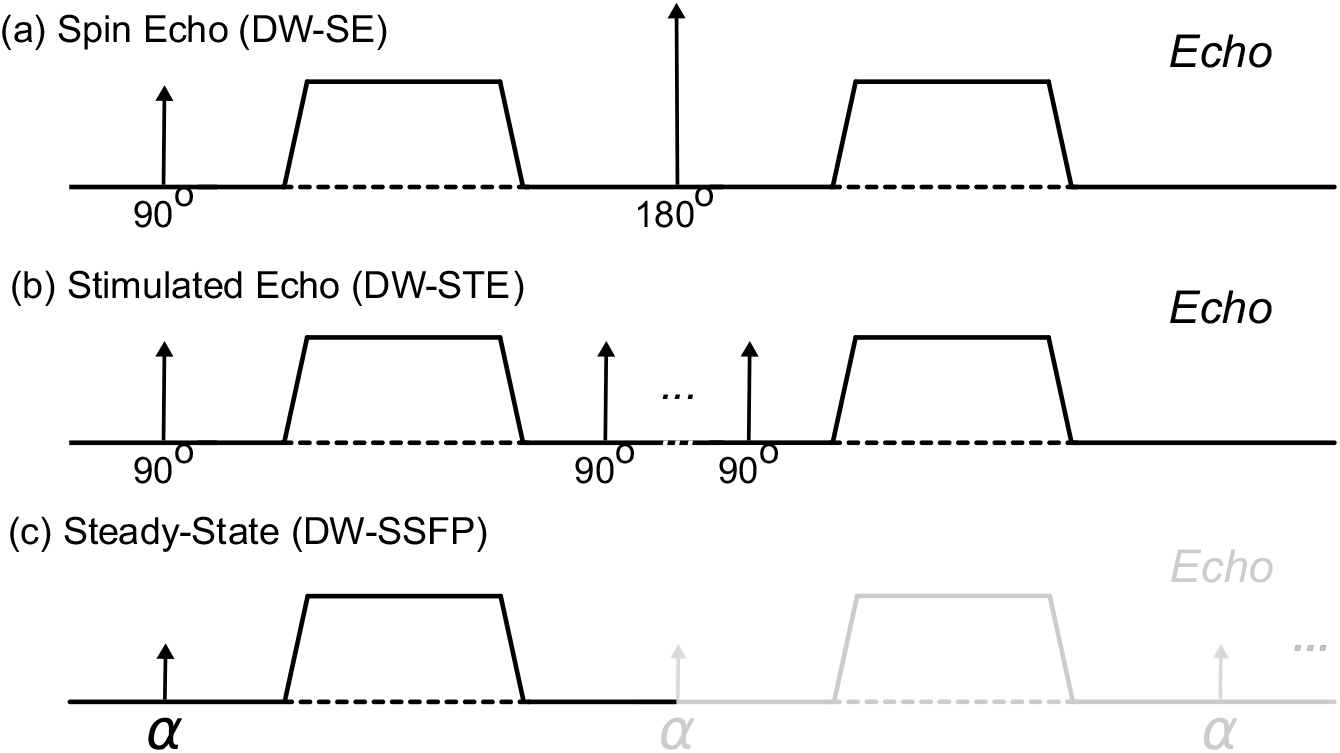
Diffusion encoding of different MRI sequences. The diffusion-weighted spin-echo (DW-SE) (a); diffusion-weighted stimulated echo (DW-STE) (b); and diffusion-weighted steady-state free precession (DW-SSFP) (c) sequence. DW-STE (b) achieves strong diffusion-weighting with reduced T_2_ signal loss by increasing diffusion sensitisation longitudinally (associated with slow T_1_ recovery). This results in increased experimental time and signal-forming mechanisms that lead to a 2-fold reduction in signal levels when compared to a DW-SE. (c) DW-SSFP consists of a single RF pulse and diffusion gradient per TR. Magnetisation persists and evolves over multiple TRs (see Theory), which can lead to high SNR-efficiency and strong diffusion weighting.

While DW-SE is the leading diffusion MRI method, the sequence has several well-established limitations. These limitations often arise from its (1) long diffusion-encoding period and (2) EPI readout^[2]^. In recent decades, a key focus of MRI methods development has been the establishment of new technologies to address these limitations. This includes the development of (1) specialised gradient systems^[3–5]^, (2) accelerated acquisition methods^[6]^ and (3) novel image-reconstruction algorithms^[6]^.

Broadly, any MRI sequence can be made sensitive to diffusion with the correct placement of encoding gradients. Different sequences have their own advantages and disadvantages, and some offer potential benefits over conventional DW-SE that warrant exploration, particularly in domains where DW-SE struggles to achieve high diffusion-sensitivity or adequate SNR. One alternative is the diffusion-weighted stimulated-echo^[7]^ (DW-STE) sequence (Figure 1b), achieving higher b-values with a typical trade-off of reduced SNR-efficiency, due primarily to a fundamental factor of two signal reduction compared to DW-SE.

In this work, I investigate Diffusion-Weighted Steady-State Free Precession^[8–11]^ (DW-SSFP) (Figure 1c), a powerful diffusion imaging sequence that has demonstrated high SNR-efficiency and strong diffusion-weighting^[12]^ with minimal image distortions (no EPI readout required). Whilst in vivo use is plagued by high motion sensitivity^[13]^, DW-SSFP has become an established method for post-mortem imaging^[14–22]^, addressing the low-diffusivity and short T_2_ environment of fixed post-mortem tissue.

A key challenge for DW-SSFP is signal interpretation. Unlike conventional diffusion MRI sequences, DW-SSFP does not have a well-defined b-value^[13,23,24]^, and complicated dependencies on both tissue relaxation properties (T_1_ and T_2_) and sequence parameters (flip angle and TR)^[13,25]^. This means that DW-SSFP does not probe a well-defined diffusion ‘regime’ in as straightforward a manner as DW-SE, limiting both interpretation and the integration of diffusion models for characterising tissue microstructure. The deviation of DW-SSFP contrast from familiar DW-SE contrast mechanisms is embodied in the deviation of the diffusion-encoding module from conventional diffusion gradient pairs: DW-SSFP contains only a single diffusion gradient in each TR (Figure 1c – black line).

In this paper, I first address the challenge of DW-SSFP signal interpretation. Building on Extended Phase Graphs (EPG)^[26]^, I establish two new representations of the DW-SSFP signal in terms of (1) conventional b-values (time-*independent* diffusion) and (2) encoding power-spectra^[27]^ (time-*dependent* diffusion). These representations build on existing DW-SSFP descriptions by incorporating the specific gradient waveform and timings experienced by distinct signal-forming pathways, facilitating translation of a DW-SSFP measurement into an interpretable diffusion regime (extending existing partition framework representations of DW-SSFP^[11,25]^). I use these representations to visualise how the investigated diffusion regime is impacted by both sequence parameters and alternative gradient waveforms (oscillating gradients), facilitating the identification of regimes where DW-SSFP may offer benefits for characterising tissue microstructure over conventional sequences.

I subsequently use these representations to establish a framework for characterising the impact of time-*independent* and -*dependent* diffusion on the measured DW-SSFP signal. Notably, existing diffusion models can be incorporated into the proposed framework without the requirement of extensive derivations. Investigations are performed considering both free Gaussian diffusion and a restriction system (cylinder of varying radius), achieving excellent agreement to existing analytical models of the DW-SSFP signal and Monte Carlo simulations. From the perspective of signal-forming mechanisms, these frameworks provide intuition into how DW-SSFP conceptually achieves higher levels of diffusion attenuation compared to a conventional DW-SE under the condition of matched diffusion encoding waveforms and timings. I apply the time-*independent* framework to experimental DW-SSFP data acquired in a whole post-mortem brain from a previously-described dataset^[28,29]^, estimating voxelwise tensor and proof of principle NODDI estimates.

Finally, I perform an investigation into the SNR-efficiency properties of DW-SSFP as a function of gradient strength, relaxation times, and encoding period (oscillating gradients). I find that whilst the theoretical SNR benefits of DW-SSFP are reduced when compared to DW-SE at increased gradient strengths (and matched encoding periods), conventional (unbalanced) DW-SSFP offers improved SNR-efficiency at short T_2 &_ long T_1_ relaxation times. Combined with its low distortion properties, DW-SSFP demonstrates considerable promise for post-mortem microstructural imaging at ultra-high field.

Software is provided to enable researchers to perform their own investigations with the proposed frameworks, including scripts to replicate many of the findings presented in this manuscript (available [here](https://github.com/BenjaminTendler/SteadyStateDiffusionMicrostructure)).

### Theory

#### Overview of DW-SSFP

The DW-SSFP sequence consists of a single RF pulse (of flip angle *α*) and diffusion gradient (with characteristic length scale *q*^−1^) in each repetition time (Figure 1c – black line)^[13]^. As with all steady-state sequences, the repetition times are short relative to the tissue T_2_ (TR typically 20 - 40 ms).

Remaining transverse magnetisation is not spoiled at the end of each TR, leading to magnetisation that experiences repeated sensitisation to RF pulses and diffusion gradients (Figure 1c – grey line).

In DW-SSFP, magnetisation accumulates diffusion contrast over several TRs, consistent with dephasing and rephasing of magnetisation due to gradient pairs in conventional diffusion MRI sequences. The resulting steady-state can be considered a superposition of multiple magnetisation components with different histories – e.g., pairs of diffusion-weighted gradients separated by a variable integer number of TRs that lead to different degrees of diffusion weighting. This creates a composite signal corresponding to a summation of these components, with non-trivial diffusion-weighted signal attenuation.

For unrestricted ‘free’ Gaussian diffusion, an analytical expression of the DW-SSFP signal exists^[30]^ (Appendix 1), analogous to, but far more complicated than, the characteristic exponential decay (*S*_0_*e*^*b*⋅*D*^) associated with conventional diffusion MRI sequences. Examination of Appendix 1 provides insights into the dependencies of DW-SSFP. Diffusion-weighted terms (*e*^…*D*^) are not separable from the remainder of the equation, leading to diffusion-attenuation additionally dependent on relaxation properties (T_1_ and T_2_), sequence flip angle (*α*), and TR (Figure 2).

**Figure 2:**
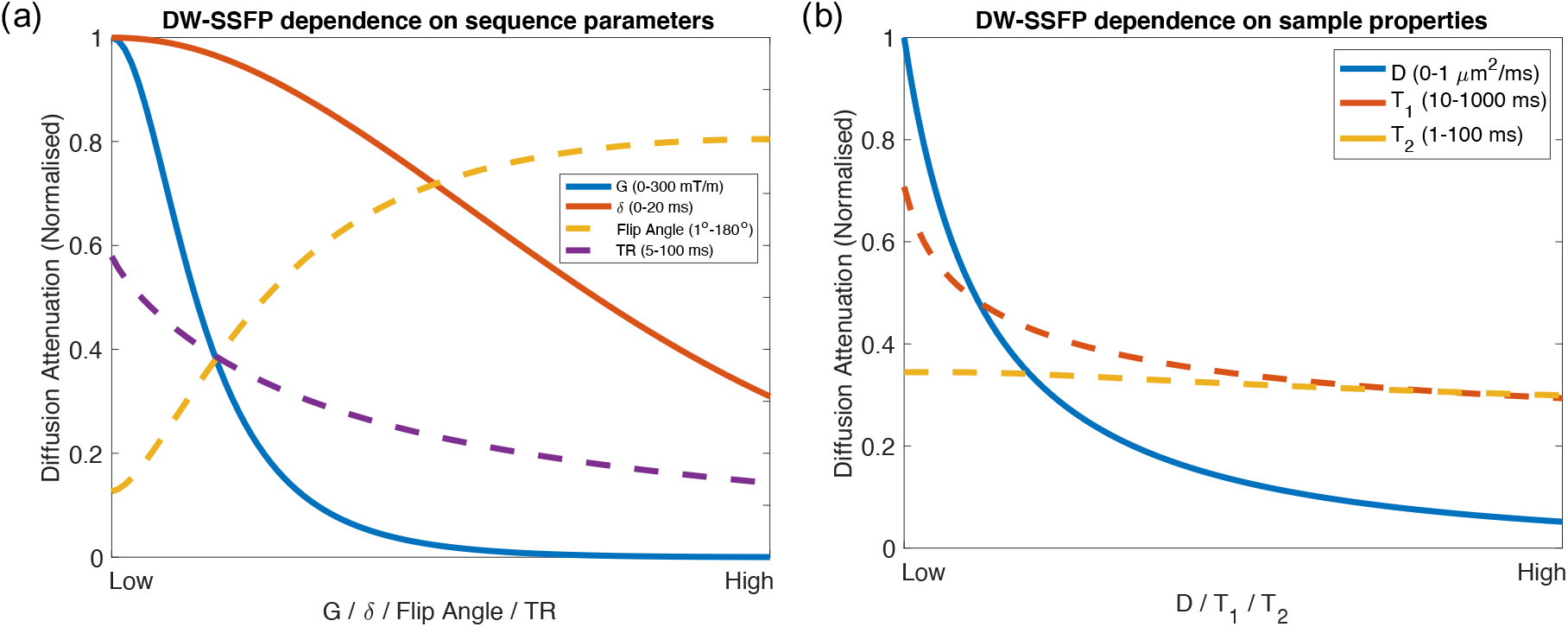
DW-SSFP diffusion attenuation. Dependence of DW-SSFP diffusion attenuation on (a) sequence parameters and (b) sample properties. Whilst conventional diffusion acquisitions are influenced by the diffusion gradient (*G*), duration (*δ*) and coefficient (*D*) (solid lines), several further parameters influence DW-SSFP diffusion attenuation (dashed lines). In DW-SSFP, decreasing the flip angle / increasing the TR corresponds to increased diffusion-weighting (a). A higher T_1_ or T_2_ also leads to increased diffusion-weighting (b). DW-SSFP signal estimated using the analytical model described in Freed et al.^[30]^ with gradient duration *δ* (Appendix 1). Sequence parameters based on post-mortem DW-SSFP investigations at 7T as described in Tendler et al.^[29]^ (defined in Methods). Note that there is no explicit definition of diffusion time (Δ) for the DW-SSFP sequence. To explore these relationships further, find associated code [here](https://github.com/BenjaminTendler/SteadyStateDiffusionMicrostructure).

The DW-SSFP sequence under free Gaussian diffusion can also be investigated using Extended Phase Graphs (EPG)^[26]^. Briefly, EPG describes a widely adopted framework for modelling the magnetisation evolution of an MRI sequence. It provides a solution to the Bloch equations^[31]^, reparameterising magnetisation as a Fourier basis with distinct phase states (*k*), defining:

- Longitudinal 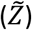 and transverse 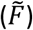 magnetisation components with different phase (*k*) states.
- RF pulses acting as a mixing operator on 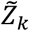 and 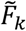 components.
- Diffusion gradients acting on 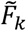 components.

EPG can characterise the estimated diffusion attenuation for an arbitrary train of RF pulses and diffusion gradients, with existing work incorporating both free Gaussian diffusion and extensions to diffusion anisotropy^[26,32]^. For readers unfamiliar with EPG signal representations, I recommend the review article by Weigel^[26]^.

#### Modelling tissue microstructure with DW-SSFP

It is well established that diffusion in tissue is not in general characterised by free Gaussian diffusion^[33]^. Several features of tissue microstructure hinder and restrict the diffusion of water, the majority reflecting the presence of membranes with very limited permeability on the timescales of diffusion MRI measurements. This has motivated the development of sophisticated models and experimental methods to relate diffusion MRI measurements to microstructural features^[33]^.

In this work, I separate diffusion processes into two distinct domains. Time-*independent* diffusion processes characterise diffusion as Gaussian, where a sequence’s b-value is sufficient to fully describe the measured diffusion attenuation. This always holds for free-diffusion, and is also valid for microstructural systems in the long-diffusion time regime^[33]^. Time-*dependent* diffusion systems incorporate non-Gaussian diffusion processes (e.g. arising from restrictions experienced by mobile spins). Here, information about the specific gradient waveform timings, *G*(*t*), are required to characterise the measured diffusion attenuation^[33–35]^.

For time-*independent* diffusion, previous DW-SSFP investigations have taken existing analytical representations of free Gaussian diffusion^[25,30]^ and extended them to incorporate models consisting of multiple Gaussian compartments including (1) tensors^[14]^, (2) ball & sticks^[14,36]^, and (3) a Gamma distribution^[24]^. Resulting estimates have been used to perform tractography^[14,15,18,19,29]^ and characterise the non-Gaussianity of tissue^[24]^. These extensions required the derivation of novel analytical forms^[14,36]^ or numerical methods^[24]^ to integrate the diffusion model. This rapidly becomes non-trivial when considering models incorporating sophisticated orientation distributions (e.g. Watson^[37]^ and Bingham^[38]^ distributions).

For time-*dependent* diffusion, it is not possible to use EPG or existing analytical models of DW-SSFP to characterise the measured signal. Specifically, whilst these approaches preserve information about gradient waveform timings within a *single* TR, they do not preserve this information across *multiple* TRs. For example, in EPG magnetisation pathways are uniquely characterised by their current phase state (*k*). This does not consider their evolution history, and pathways that ultimately end up in the same state may have experienced a different effective gradient waveform, 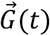, over time (Figure 3). Existing analytical DW-SSFP models are similarly impacted.

**Figure 3:**
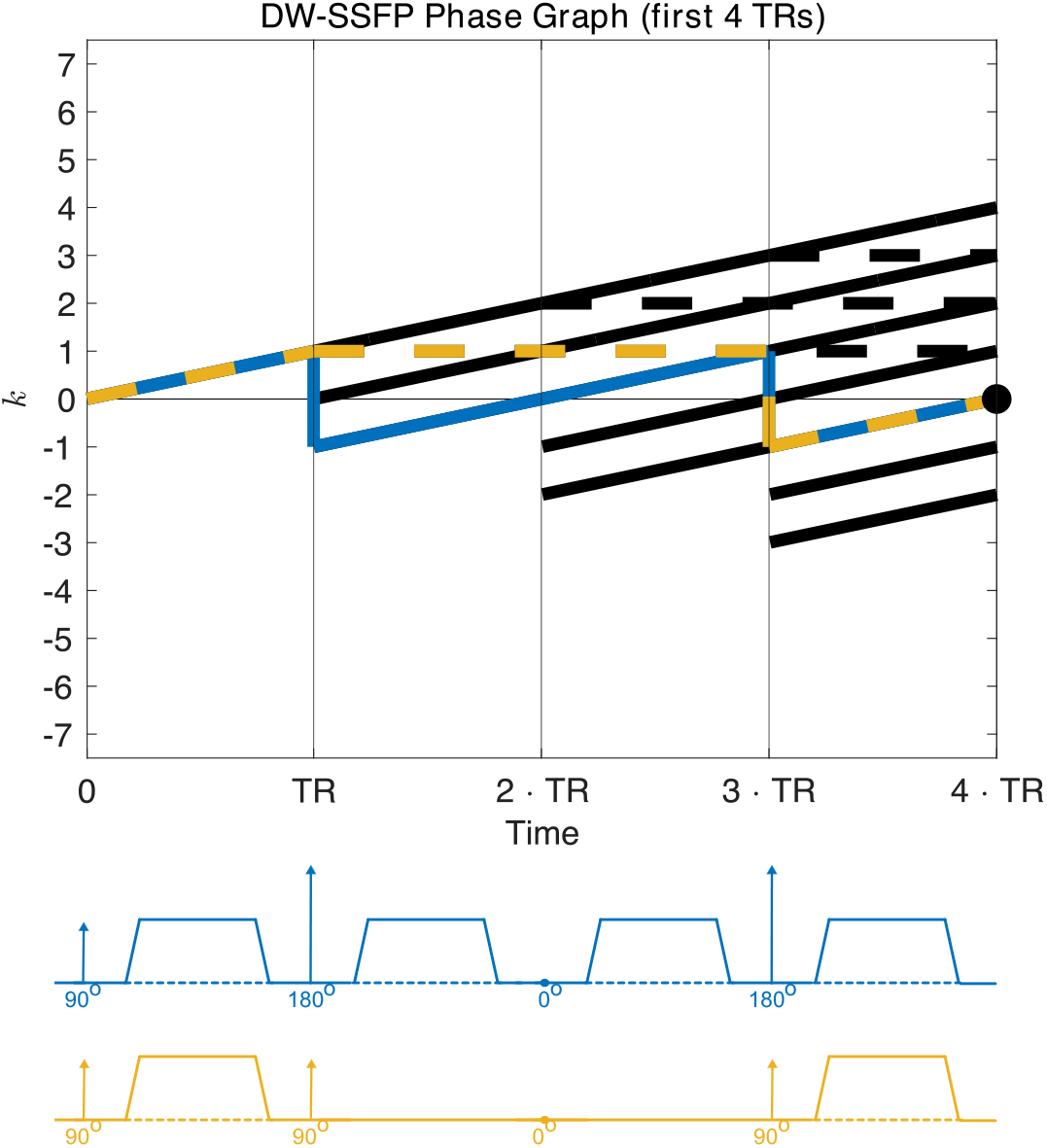
EPG representation of the DW-SSFP sequence. Using a phase graph representation^[26]^ of DW-SSFP (first four TRs), we can identify several pathways that contribute to the measured signal at the 4^th^ TR. The two highlighted pathways (blue & yellow) have an identical phase state at the 4^th^ TR (black dot), but distinct diffusion encoding trajectories (corresponding to double diffusion encoding^[34] &^ a stimulated echo^[7]^ respectively). The full gradient waveforms of these pathways must be considered to accurately estimate the DW-SSFP signal arising from a time-*dependent* microstructural system. Notably, even pathways with identical b-values but different gradient trajectories can lead to different signal estimates when considering time-*dependent* diffusion phenomena. The proposed framework distinguishes these pathways based on unique evolution histories.

Previous investigations have utilised an approximation regime of the DW-SSFP signal to demonstrate sensitivity to time-*dependent* restrictions^[23,24]^. Whilst this sensitivity could be more accurately explored with existing simulation methods (e.g. Monte Carlo simulations), to date no such investigations have been performed, and would benefit from complementary investigation methods that facilitate interpretation of the measured signal.

Below, I describe an approach to address the above challenges, introducing a DW-SSFP framework that (i) can estimate diffusion attenuation arising from time-*independent* and -*dependent* diffusion systems, (ii) embeds existing diffusion models without the requirement of extensive derivations; (iii) translates defined DW-SSFP parameters into an interpretable diffusion regime; and (iv) can investigate the impact of alternative gradient waveforms on the DW-SSFP signal.

#### Overview

The proposed framework builds on EPG to distinguish pathways that have an identical phase (*k*) state, but different evolution histories. It can be alternatively interpreted as an extension to existing partition framework^[11,25]^ representations of DW-SSFP.

Building on the notation in Weigel^[26]^, I characterise the measured DW-SSFP signal as:

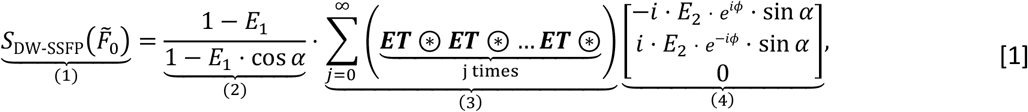

consisting of:

1. The measured DW-SSFP signal (sum of all 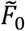 pathways)
2. Steady-state longitudinal magnetisation
3. Application of the RF-excitation & relaxation EPG operator^[26]^
4. Initialised magnetisation vector in the transverse plane 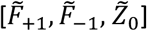

where 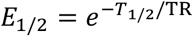, ⊛ defines a pathway expansion operator (described below) and^[26]^:

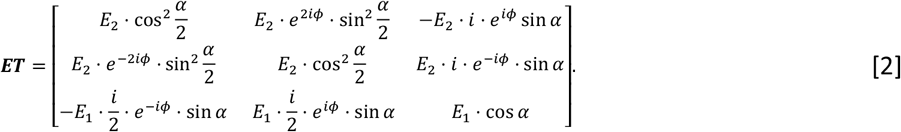

I distinguish pathways based on their evolution history via the pathway expansion operator:

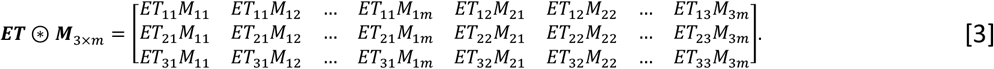

⊛in Eq. [1] can be interpreted as separating each magnetisation component mixed by the ***ET*** operator. Specifically, whereas conventional EPG sums magnetisation components with identical 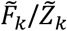 states (with each TR adding an additional set of states) the proposed framework increases the number of states threefold per TR. This leads to a matrix of dimensions 3x3^m^, where the top/middle and bottom rows correspond to 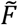 and 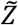 states respectively.

I identify signal-contributing 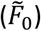 magnetisation by separately tracking the phase state of each pathway via a shift operator 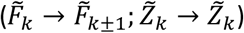, where *k* can be positive or negative. The phase state evolution (*k*(TR_1_) → *k*(TR_2_) → *k*(TR_3_) → …) of each signal-forming pathway is stored and used to synthetise its experienced gradient waveform, 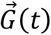 (Figure 3). I motivate Eqs. [1]-[3] with two application examples in Supporting Information.

To account for magnetisation components that are insensitive to diffusion, Eq. [1] initialises all pathways in the transverse plane (4). This initialisation builds on the principle that equilibrium longitudinal magnetisation 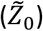 experiences no gradient and is therefore insensitive to the effects of diffusion, meaning it can be modelled as a scalar constant (2). This captures the effects of all ‘delayed’ pathways (i.e. pathways that persist for several TRs in 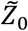 prior to initial excitation into the transverse plane) and 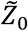 pathways arising from longitudinal magnetisation recovery.

#### Pathway identification and storage

In practice, the number of pathways arising from Eq. [1] scales as 3^n^ (n = number of TRs). Here, I mitigate the challenge of dimensionality by exploiting degeneracies in pathway evolution. For example, Figure 3 displays two pathways (in blue and yellow) that have distinct diffusion properties and signal amplitudes, but identical phase states at the fourth TR. Following the fourth TR the magnetisation associated with these pathways will evolve in an identical manner, only distinguished by their initial signal amplitude and effective gradient waveform during the first four TRs.

Using this concept and the initialisation in (Eq. [1]), we can represent the evolution of any signal-forming pathway via:

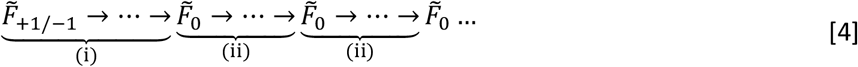

Specifically, we can describe a pathway using two distinct components. The first component (i) describes the evolution of the pathway from an initial 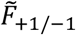 state to a signal-forming 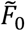 state. The second component (ii) describes the evolution of the pathway from the 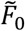 state to a further signal-forming 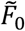 state. Longer pathways can be described by recursively appending components that are consistent with the definition of (ii).

Using this approach, I condense the information required to simulate all signal-forming pathway into two dictionaries. These dictionaries contain the signal amplitude and gradient waveform of all pathways evolving via (i) or (ii). We can estimate the properties of an arbitrary signal forming pathway by combining an entry in dictionary (i) with one or more entries in dictionary (ii), multiplying signal amplitudes and appending gradient waveforms. I display the gradient waveforms associated with three example dictionary components in Figure 4 (left column).

**Figure 4:**
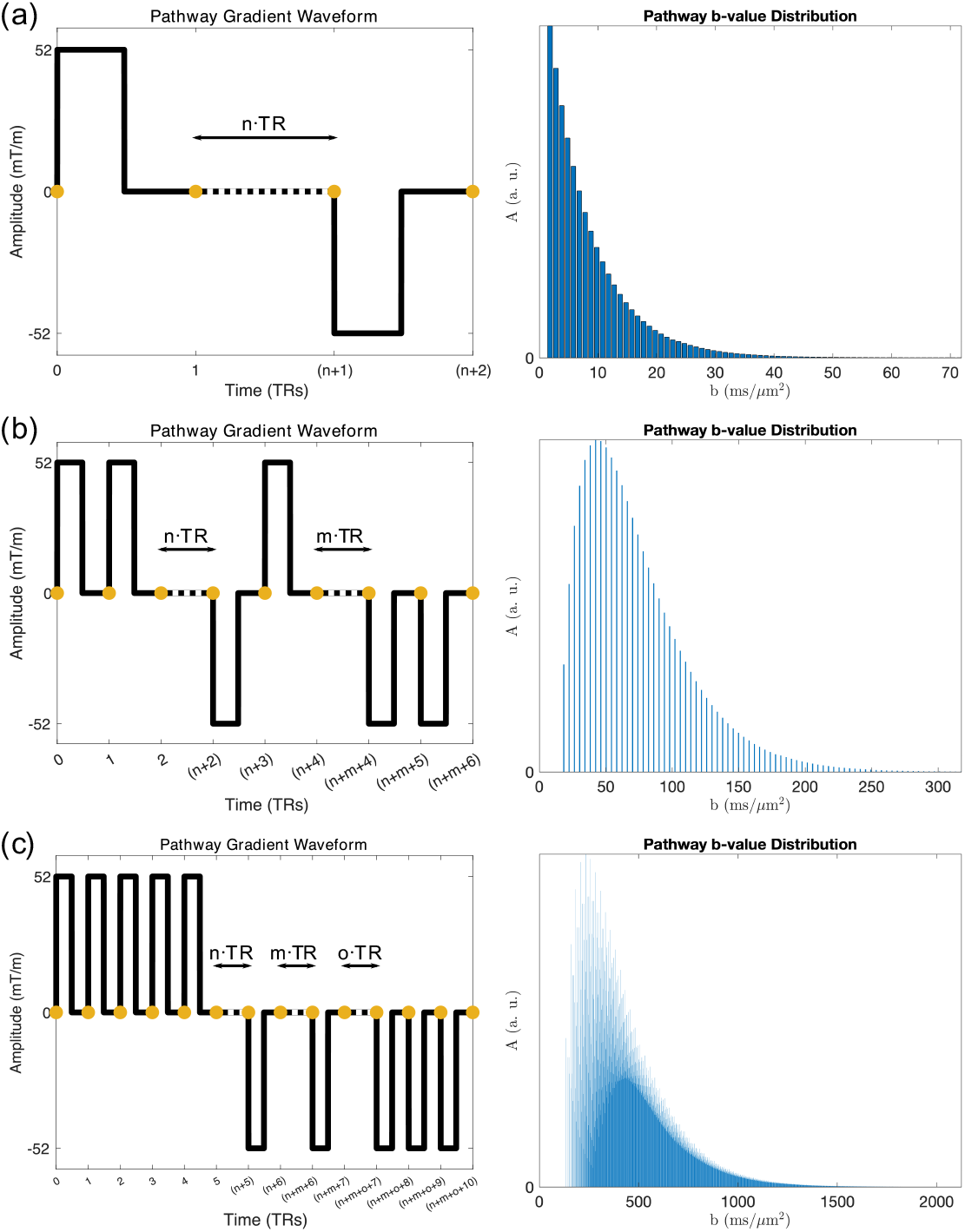
Example b-value distributions associated with dictionary components. Gradient waveforms experienced by three example dictionary entries (a-c, left) and their corresponding b-value distributions (a-c, right). (a) is the dictionary entry for all stimulated echoes. Here, the b-value distribution represents stimulated-echoes persisting for 1 to n TRs longitudinally. Dictionary entries with more than one longitudinal period (b & c) lead to more complicated distributions, arising in part due to different combinations of longitudinal period durations (i.e different *n, m, o*, …) achieving identical b-values, and distinct phase states associated with each longitudinal period. Properties of the three dictionary entries are provided in Supporting Information Table S1. Yellow circles correspond to TR timings.

The dictionaries are further condensed by representing pathways that have identical gradient moments but experience a different number of longitudinal TRs 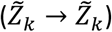 as a single entry. For example, Figure 4a (left) represents the gradient waveform of a single dictionary entry associated with all stimulated-echo pathways. This step is performed as we can exploit simple mathematical relationships between these pathways to rapidly estimate their properties, as described in the next section.

To prevent dictionaries from having an infinite number of entries, we can define simple amplitude thresholds and limits on pathway durations. For the default parameters used in this manuscript (see Methods), dictionaries (i) and (ii) contained 23,474 and 46,948 entries respectively. They were generated in under 20 seconds on a personal laptop, requiring ∼ 31 MB.

#### Signal Representation 1: Time-independent diffusion

The first representation characterises the DW-SSFP signal in the time-*independent* regime by reparameterising the measured DW-SSFP signal as a b-value distribution. I achieve this by first estimating the b-value distribution associated with each dictionary component. For example, Figure 4a (right) displays the b-value distribution associated with all stimulated-echo pathways.

Using the stimulated-echo entry as an example (Figure 4a), I synthesise the b-value distribution by estimating the signal-amplitude and b-value associated with each instance of a different number of TRs between the two diffusion gradients via:

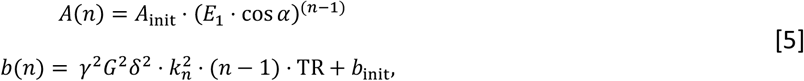

where *n* defines an integer number of TRs between the two diffusion gradients (Figure 4a), *k*_*n*_ is the integer phase state associated with the longitudinal period of the stimulated-echo, *A*_init_ is the signal amplitude the shortest (*n* = 1) stimulated-echo, and *b*_init_ is the b-value associated with the shortest stimulated-echo (Supporting Information Table S1). Each bar in Figure 4a (right) corresponds to the signal amplitude and b-value associated with a different *n*.

We can generalise the above equation for arbitrary dictionary entries with several longitudinal periods (corresponding indices *n, m, o*, …) by defining:

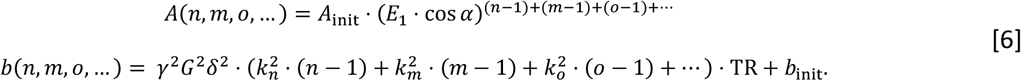

Modulation of the b-value by 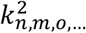 reflects the increased q-value associated with longitudinal periods that have experienced repeat sensitisation to a diffusion gradient (*q* = *γGδ* · *k*_*n,m,o*,…_).

Figure 4 displays the b-value distribution of three example dictionary components, with their properties defined in Supporting Information Table S1. As combinations of different longitudinal period durations (i.e different *n, m, o*, …) can share the same b-value, we can accurately represent the b-value distribution with reduced dimensionality.

Building on this approach, we can rapidly synthesise b-value distributions for different dictionary entries by exploiting simple relationships between them. For example, synthesising the b-value distribution of a dictionary entry with two longitudinal periods (example in Figure 4b) requires estimation of the b-value and signal amplitude corresponding to *n* = 1, *m* = 1 → ∞; *n* = 2, *m* = 1 → ∞; *n* = 3, *m* = 1 → ∞ etc. This description is analogous to a convolution (*) of two dictionary entries with one longitudinal period (Supporting Information Figure S1), defining:

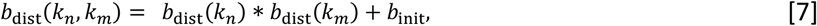

where *b*_dist_(*k*_*n*_, *k*_*m*_) is a b-value distribution corresponding to a dictionary component with two longitudinal periods (phase states *k*_*n*_ and *k*_*m*_), *b*_dist_(*k*_*n*_) and *b*_dist_(*k*_*m*_) are b-value distributions corresponding to a dictionary component with one longitudinal period (phase state *k*_P_ or *k*_T_), and *b*_init_ is the b-value of the shortest gradient waveform associated with the dictionary entry with two longitudinal periods (*n* = *m* = 1). The above relationship can be generalised for dictionary entries with several longitudinal periods via:

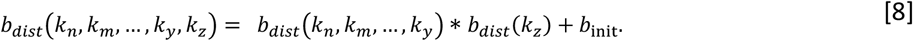

The above relationships state that the shape of the b-value distribution associated with a dictionary entry is uniquely defined by (1) the number of distinct longitudinal periods (*n*_*long*_) and (2) the phase state associated with each longitudinal period (*k*_*n,m,o*,…_). The only difference in the b-value distribution of dictionary entries with identical *n*_init_ and *k*_*n,m,o*,…_ is associated with their b-value offset *b*_NQNR_ (Supporting Information Figure S2). The 23,474 and 46,948 entries in dictionaries (i) and (ii) are associated with 680 unique b-value distribution shapes.

Once we have identified the b-value distribution associated with each entry in dictionaries (i) and (ii) (Eq. [4] and surrounding text), we can sum them together to create two b-value distributions, 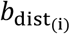 and 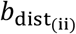. We can generate the final b-value distribution via a similar recursive convolution operation:

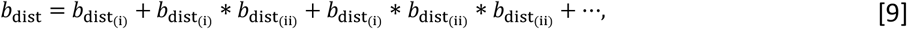

where the different components in the above equation correspond to pathways that evolve via 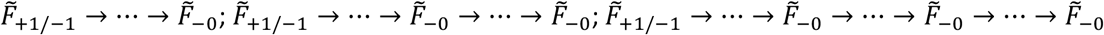 etc.

Figure 5 displays the final b-value distribution associated with the default investigation parameters in this manuscript (see Methods). It reparameterises the DW-SSFP signal as the weighted-sum of signal fractions with different b-values. It represents the information of over 10^18^ pathways, and was synthesised in under 15 seconds on a personal laptop. Note that pathways can have positive or negative signal amplitudes (Supporting Information Figure S3).

**Figure 5:**
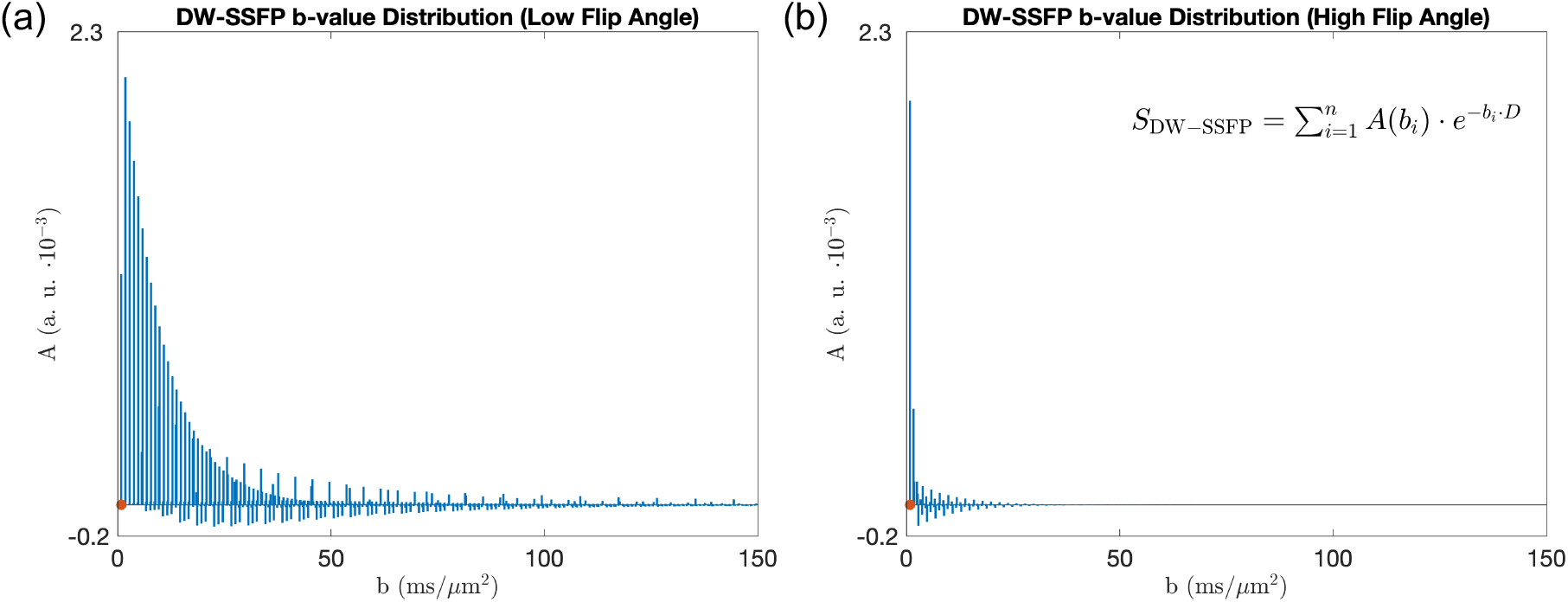
DW-SSFP b-value representation. By estimating the b-value and amplitude of each signal-forming pathway, the measured DW-SSFP signal can be represented as the weighted sum of signal fractions with different b-values (*A* defined in Eq. [10]). Here I visualise this representation for DW-SSFP data acquired at a (a) low (20 °) and (b) high (160 °) flip angle, providing insights into how changing a DW-SSFP parameter translates into a given diffusion regime. We can subsequently estimate the measured DW-SSFP signal by summing over the b-value range (x-axis), scaling the amplitudes with an appropriate diffusion model (Eq. [10] - inset equation displays the signal model for free Gaussian diffusion). The Orange dots indicate the equivalent DW-SE b-value assuming identical gradient amplitudes and timings (setting Δ = TR), equal to 0.836 ms/μm^2^. Parameters based on post-mortem DW-SSFP investigations at 7T as described in Tendler et al.^[29]^ (defined in Methods). To explore how changing parameters influences the distributions, find the associated code here.

This representation provides insight into the diffusion regimes investigated by DW-SSFP. By changing the sequence parameters or sample properties, we observe a change to the relative-weighting of pathways with different b-values. Here, lower DW-SSFP flip angles corresponds to an increased signal fraction at higher b-values, lead to greater diffusion-weighting of the DW-SSFP signal (Figure 2). It also demonstrates how DW-SSFP achieves strong diffusion-weighting, with the majority of signal arising from b-values that exceed those arising from a DW-SE with identical diffusion-encoding gradient and timings.

A powerful aspect of this representation is that it also enables us to directly translate existing time-*independent* models into DW-SSFP without deriving novel analytical forms. Specifically, we can use the b-value distribution to represent the measured DW-SSFP signal as:

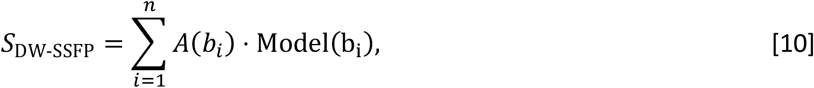

where *A*(*b*_*i*_) is the signal amplitude at each b-value (Figure 5), *n* is the number of b-values in the distribution and Model(*b*_*i*_) defines any time-*independent* model. For example, for free Gaussian diffusion:

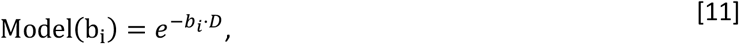

for a Tensor:

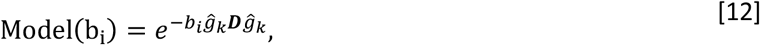

for NODDI^[39]^

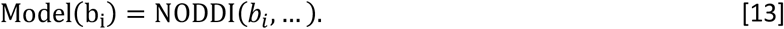

This is particularly helpful in DW-SSFP due to the complicated nature of its signal on both tissue and sequence parameters. When considering free Gaussian diffusion, Eq. [10] leads to the same result as conventional EPG or an analytical DW-SSFP model (Appendix 1).

Unlike previous partition framework implementations utilising the two-transverse^[23,25]^ or four-transverse^[40]^ period approximations of DW-SSFP, a key advantage of the proposed time-*independent* pathway framework is that it can incorporate higher-order signal components to accurately represent the DW-SSFP signal. This is particularly valuable when considering regimes where the T_2_ >> TR (Supporting Information Figure S4).

#### Signal Representation 2: Time-dependent diffusion

In this work I model time-*dependent* diffusion using the Gaussian Phase Approximation^[27,41,42]^ (GPA). In contrast to a waveform’s b-value (which only considers the total gradient integral), the GPA explicitly incorporates information about the timings of the gradient waveform via the evolution of the q-vector, *q*(*t*) = *γ*∫ *G*(*t*)*dt*. It reparameterises the estimated diffusion attenuation as a function of the encoding power applied at different temporal frequencies of the diffusion spectrum (Fourier transform of the velocity-autocorrelation function^[43,44]^) of the underlying microstructural system.

Using a similar approach to the previous section, I model the DW-SSFP signal via the GPA as:

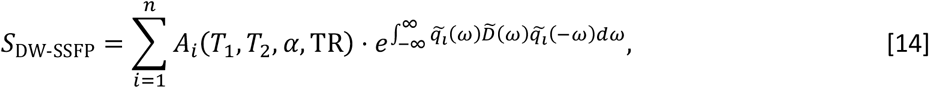

where *A*_*i*_(*T*_1_, *T*_2_, *α*, TR) is the amplitude of a signal forming pathway, 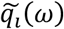 is the Fourier transform of the pathway’s q-vector, *n* is the number of pathways, and 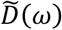 is the diffusion spectrum of the microstructural system.

Figure 6 reparameterises the DW-SSFP signal as a as density plot of gradient power-spectra. Like the b-value representation (Figure 5), by changing DW-SFP sequence parameters or sample properties we observe a change the relative amplitude and width of 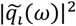 (Supporting Information Figure S5).

**Figure 6:**
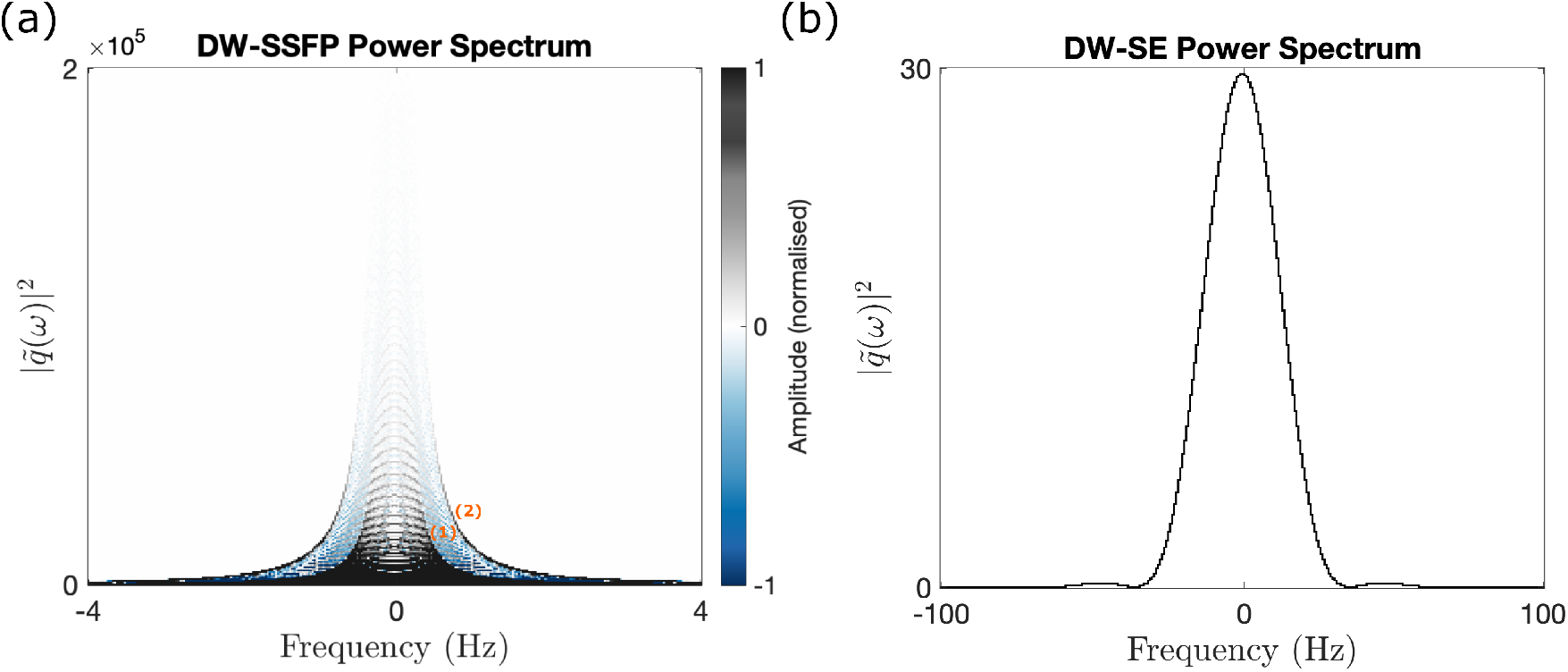
DW-SSFP power spectrum representation. (a) By estimating 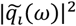 and the amplitude of each signal-forming pathway, the measured DW-SSFP signal is represented as a power spectrum density plot. Here, image colour represents the signal amplitude contributed to a given regime, with DW-SSFP pathways associated with both positive and negative amplitudes (Supporting Information Figure S3). (b) displays the equivalent power spectrum for a DW-SE sequence using identical gradient timings (setting Δ = TR). For DW-SSFP, 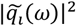 spans a narrower frequency range with substantially higher amplitudes, where both x and y axes in (a) and (b) span different extents. ‘Contours’ in (a) (labelled 1 & 2) correspond to pathways experiencing two and four periods in the transverse plane. For equivalent spectrums at different flip angles, see Supporting Information Figure S5. (a) displays 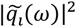 for pathways that experience up to four periods in the transverse plane. Parameters based on post-mortem DW-SSFP investigations at 7T as described in Tendler et al.^[29]^ (defined in Methods). To explore how changing parameters influences the power spectrum, find the associated code [here](https://github.com/BenjaminTendler/SteadyStateDiffusionMicrostructure).

This representation demonstrates that the DW-SSFP signal is typically strongly concentrated in a narrow range of the frequency spectrum when compared to a conventional DW-SE.

Unlike the highly compressible signal representation for time-*independent* diffusion, there is little degeneracy of 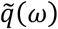 for pathways that contribute to the measured DW-SSFP signal. For the time-*dependent* diffusion representation I use the dictionaries (Theory: *Pathway identification and storage*) to explicitly generate the full gradient waveform and signal amplitude of *each* individual signal-forming pathway. I achieve this by recursively appending the gradient waveforms from the first 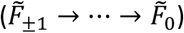 and second 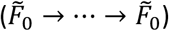 dictionaries and multiplying their signal amplitudes. I identify the location of each longitudinal period (Figure 4 – dashed lines) and extend each gradient waveform for different periods in the longitudinal plane, scaling the pathway signal amplitudes by (*E*_1_ ⋅ *cos α*)^(*n*−1) + (*m*−1) + (*o*−1) +…^ (Eq. [6]).

A key challenge of the time-*dependent* pathway framework is the requirement to explicitly incorporate the full gradient waveform and signal amplitude of each pathway separately, leading to long computational times. To address this, in this work I limited investigations to pathways that persist for up to four TRs in the transverse plane (four transverse-period approximation). This approximation corresponds to ∼99% of the measured signal for the investigated parameter regime (Supporting Information Figure S6), corresponding to ∼275,000 pathways.

#### Alternative Gradient Waveforms

We can use the above frameworks to investigate the impact of alternative gradient waveforms on the DW-SSFP signal by explicitly integrating the waveform shape into the investigation. This can be achieved without additional coding or analytical derivations.

Figure 7 displays the predicted DW-SSFP b-value distribution and power-spectrum with oscillating gradients (sinusoidal waveform and parameters based on Aggarwal et al.^[45]^). The b-value distribution (Figure 7b) arising from the time-in*dependent* framework provides an efficient description of the DW-SSFP signal, condensing the entire measurement into a few discrete signal components with different b-values (scalar multiples of the b-value per TR). The distribution corresponds to the fractions of signal that experience 2 ⋅ *n* TRs in the transverse plane (integer *n*).

**Figure 7:**
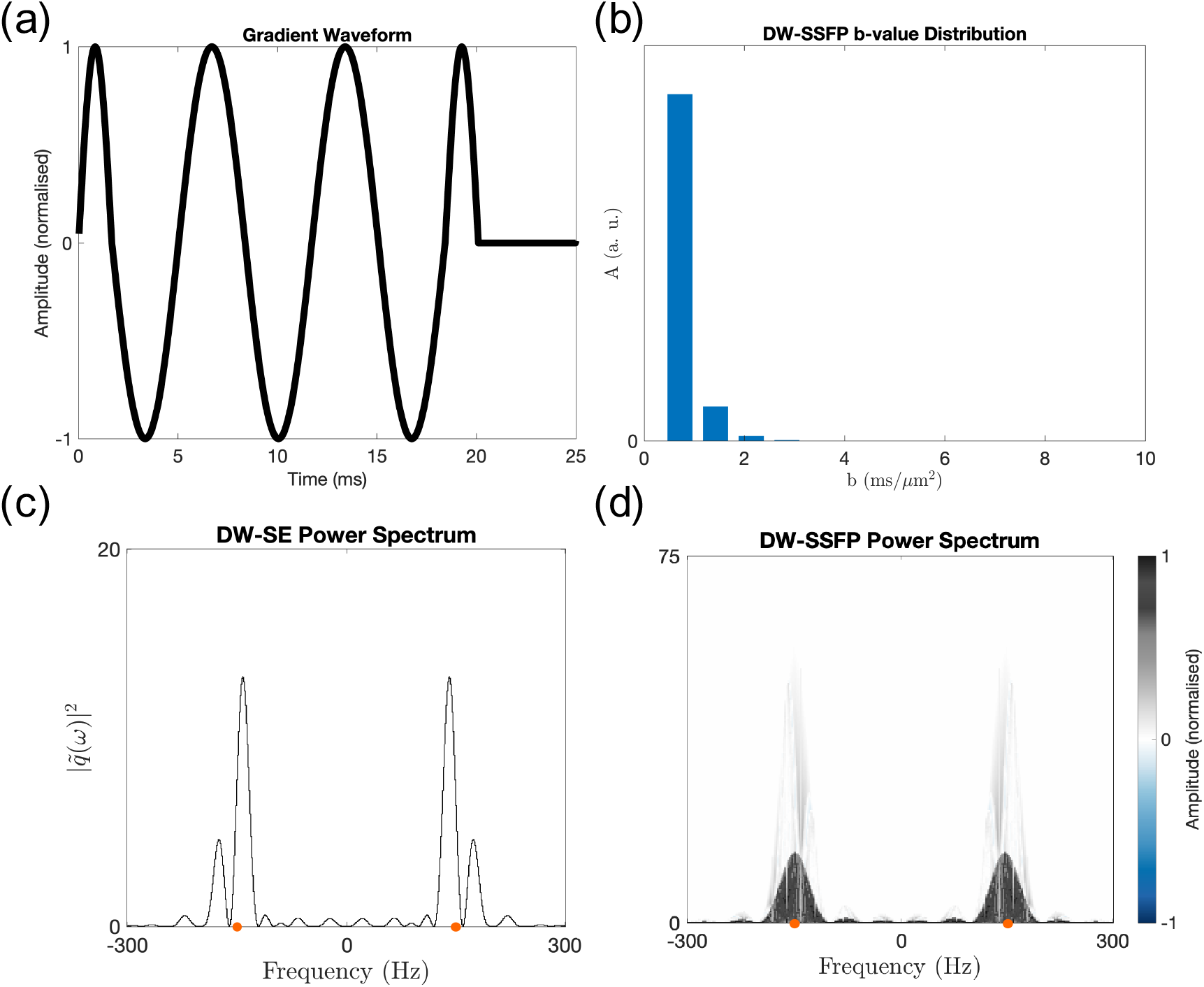
DW-SSFP with oscillating gradients. Using the balanced sinusoidal gradient waveform in (a) (equivalent to Figure 1d in Aggarwal et al.^[45]^ - oscillation frequency = 150 Hz), we can characterise the DW-SSFP signal with oscillating gradients as a b-value distribution (b). Here, each peak corresponds a 2 ⋅ *n* multiple of the waveforms b-value (*b* = 0.35 ms/μm^2^). (c) represents the power-spectrum for a DW-SE using the gradient waveform in (a) (analogous to Figure 1e in Aggarwal et al.^[45]^), with peaks centred around ±150 Hz (orange dots). By estimating 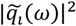 and the amplitude of each signal-forming pathway, the DW-SSFP power spectrum density plot (d) displays similar characteristics, with peak amplitudes centred around ±150 Hz (orange dots). This shows that DW-SSFP is capable of probing different frequency components of the diffusion spectrum, as reflected by deviation of central peak of the power spectrum plot away from 0 Hz. Parameters based on Aggarwal et al.^[45]^ (see Methods). (d) displays 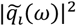 for pathways that experience up to four periods in the transverse plane. To explore how changing parameters influences the distributions with oscillating gradients, find the associated code [here](https://github.com/BenjaminTendler/SteadyStateDiffusionMicrostructure).

The time-*dependent* pathway framework demonstrates that DW-SSFP with integrated oscillating gradients can probe different frequency components of the diffusion spectrum (Figures 7c and d), here centred around the oscillation frequency of the sinusoidal gradient waveform (*f* = 150 Hz). The addition of oscillating gradients into the DW-SSFP sequence is therefore analogous to the implementation of oscillating gradients for conventional diffusion MRI sequences.

Note that the DW-SSFP sequence incorporates an unbalanced gradient per TR to prevent banding artefacts arising from off-resonance effects. To address this for the balanced oscillating gradients waveform presented here, we can introduce an independent, small unbalanced gradient; slightly unbalance the oscillating waveform^[46]^; or add a small vertical offset between positive and negative gradient lobes. A sufficiently small, unbalanced gradient typically leads to negligible diffusion-weighting and can be integrated into the proposed framework.

#### Accuracy of DW-SSFP approximations

The b-value distributions arising from the time-in*dependent* framework with oscillating gradients (Figure 7b) can alternatively be used to assess the accuracy of DW-SSFP approximations. Specifically, a commonly used alternative model for characterising the DW-SSFP signal is the ‘two-transverse’ period approximation^[23,25]^, only considering pathways that persist for up to two TRs in the transverse plane. The accuracy of this approximation is reflected in the difference between the left-most component of the b-value distribution with oscillating gradients (Figure 7b, reflecting the total signal from all pathways experiencing two transverse TRs), and the remaining signal from all other components. This approach can be extended for alternative simplified signal models (e.g. four-transverse period approximation^[40]^).

The b-value distributions can also be used to provide insights into which parameter regimes approximation models are most precise. For example, the b-value distributions demonstrate that the two-transverse period approximation is *least* accurate at very low and very high flip angles (Supporting Information Figure S7). This reflects the contribution of stimulated-echo pathways that persist for several periods in the transverse plane (very low flip angles) and spin-echo pathways that experience repeated pairs of dephasing and rephasing (very high flip angles).

## Results

Figure 8a estimates the DW-SSFP signal for free Gaussian diffusion as a function of flip angle using (1) the proposed time-*independent* pathway framework, (2) an analytical DW-SSFP model^[30]^ (Appendix 1) and (3) Monte Carlo simulations. Results demonstrate excellent agreement between the three approaches, with conventional DW-SSFP demonstrating an expected increase in diffusion attenuation with reduced flip angle^[24]^ (Figure 5).

**Figure 8:**
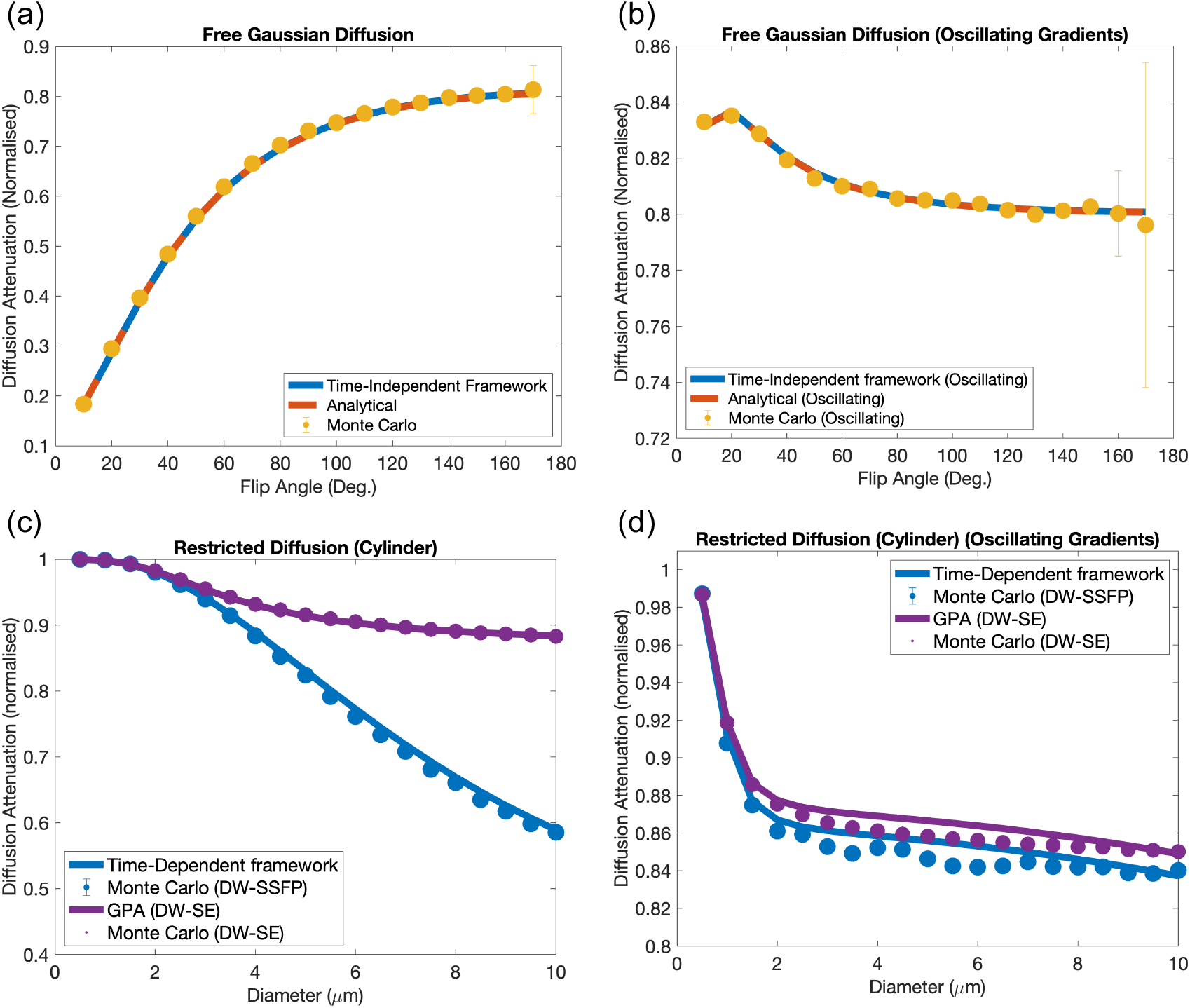
DW-SSFP signal estimation. The proposed time-*independent* framework demonstrates excellent agreement with analytical DW-SSFP signal models and complementary Monte Carlo simulations across a range of flip angles for (a) conventional (unbalanced diffusion gradient) and (b) oscillating DW-SSFP sequences. The time-*dependent* framework characterising diffusion in a single cylinder of varying radius gives good agreement to Monte Carlo simulations for conventional DW-SSFP (c), conceptually achieving higher diffusion attenuation compared to a DW-SE with matched diffusion encoding waveforms and timings. When considering oscillating gradients (d), the time-*dependent* framework gives similar accuracy to Monte Carlo simulations when compared to the DW-SE, with DW-SSFP predicting a small increase in signal attenuation. Parameters based on Tendler et al.^[29]^ for conventional gradients (see Methods) and Aggarwal et al.^[45]^ for oscillating gradients (see Methods). Comparison model of restricted diffusion for the DW-SE sequence implemented based on the Gaussian Phase Approximation^[27,41,42]^ using the proposed time-*dependent* framework. Monte Carlo simulation error bars (defined as the signal standard deviation from the final 10 TRs) are too small to be visualised for most datapoints. Note that the y axes span different extents in each plot. To explore how changing parameters influences these relationships, find the associated code [here](https://github.com/BenjaminTendler/SteadyStateDiffusionMicrostructure).

Figure 8b repeats the investigation with oscillating gradients, finding that the estimated diffusion attenuation is relatively insensitive to changes in flip angle. Here, increasing the flip angle initially leads to a reduction in diffusion attenuation, with a subsequent rise in attenuation at higher flip angles. This can be interpreted by looking at the evolution of the b-value distributions (Supporting Information Figure S7). Specifically, the relative fraction of signal that experiences 2 ⋅ n ⋅ TRs in the transverse plane is almost identical at very low and very high flip angles, despite differences in the pathways that predominantly contribute to the measured signal (see Theory: *Accuracy of DW-SSFP approximations*). Excellent agreement is again found between the three investigative approaches.

Figure 8c investigates a restricted cylinder system with varying radius, comparing the proposed time-*dependent* pathway framework to Monte Carlo simulations, with good agreement between the two approaches. Here, the average difference in attenuation between the proposed time-*dependent* framework and Monte Carlo simulations is 0.9% for the investigated parameters with DW-SSFP (0.09% for the DW-SE comparison). Comparison to a DW-SE sequence with matched gradient waveforms and timings demonstrates that DW-SSFP conceptually achieves higher levels of diffusion attenuation for the same restriction system.

Figure 8d displays a similar investigation with oscillating gradients. Here, DW-SSFP achieves a small conceptual increase in diffusion attenuation compared to a DW-SE sequence. Here, the average difference in attenuation between the proposed time-*dependent* framework and Monte Carlo simulations is 0.7% for the investigated parameters (0.5% for the DW-SE comparison).

Figures 9 a & b compare a fitted diffusion tensor derived from experimental DW-SSFP MRI data acquired in a post-mortem human brain (time-*independent* pathway framework) and an analytical DW-SSFP model^[30]^ with integrated tensor (Appendix 3). Voxelwise analysis (Figure 9c) demonstrates excellent agreement between the two approaches (Pearson correlation coefficient R=0.9992). Figure 9d displays proof of principle estimates of the intracellular volume fraction (*f*_*intra*_) and orientation dispersion (*OD*) by incorporating the NODDI model into the time-*independent* pathway framework. The resulting *f*_*intra*_ estimates demonstrate a consistently higher intracellular volume fraction within white matter. The *OD* estimates display limited contrast between white matter and grey matter in the cerebrum, with the greatest distinction within the cerebellum.

**Figure 9:**
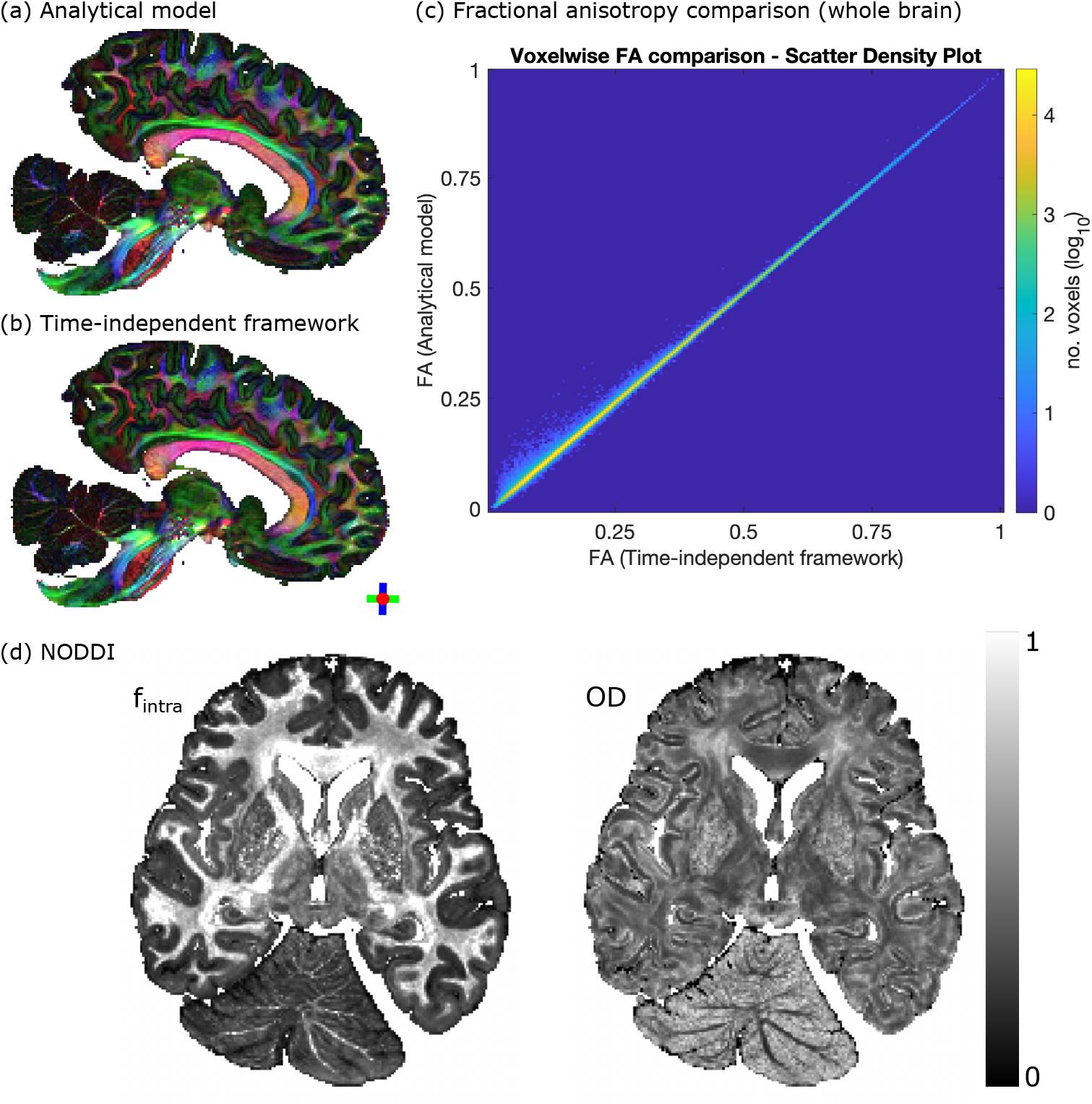
Experimental analysis. Comparison of tensor estimates derived from experimental post-mortem DW-SSFP data (whole human brain) based on the analytical DW-SSFP model^[30]^ (Appendix 3) and proposed time-*independent* pathway framework. (a & b) Fractional anisotropy (FA) modulated principal diffusion direction maps give excellent spatial agreement. (c) Scatter density plot of voxelwise FA estimates (whole brain) demonstrates the accuracy of the proposed time-*independent* pathway framework (Pearson correlation coefficient R = 0.9992). (d) Post-mortem NODDI estimates of intracellular volume fraction (*f*_*intra*_) and orientation dispersion (*OD*) derived using the time-*independent* pathway framework. Details of experimental acquisition & processing provided in the Methods.

Finally, Figures 10a-c compares the theoretical SNR-efficiency properties of DW-SSFP, DW-SE and DW-STE as a function of gradient strength (Figure 10a), T_1_ (Figure 10b) and T_2_ (Figure 10c) for matched attenuation levels (corresponding to *b* = 10 ms/μm^2^ for DW-SE). The relative SNR benefits of DW-SSFP reduce as a function of gradient strength when compared to the DW-SE (Figure 10a), with DW-SE estimating improved SNR-efficiency at G > 420 mT/m in the investigated parameter regime. When considering the maximum investigated gradient strength (G = 1000 mT/m), DW-SSFP predicts improved SNR-efficiency in tissues with long T_1_ and short T_2_ (Figures 10b and c), the relaxation conditions associated with ultra-high field. Combined with distortion-free outputs (DW-SSFP does not require an EPI readout), DW-SSFP may offer considerable benefits for ultra-high field microstructural investigations.

**Figure 10:**
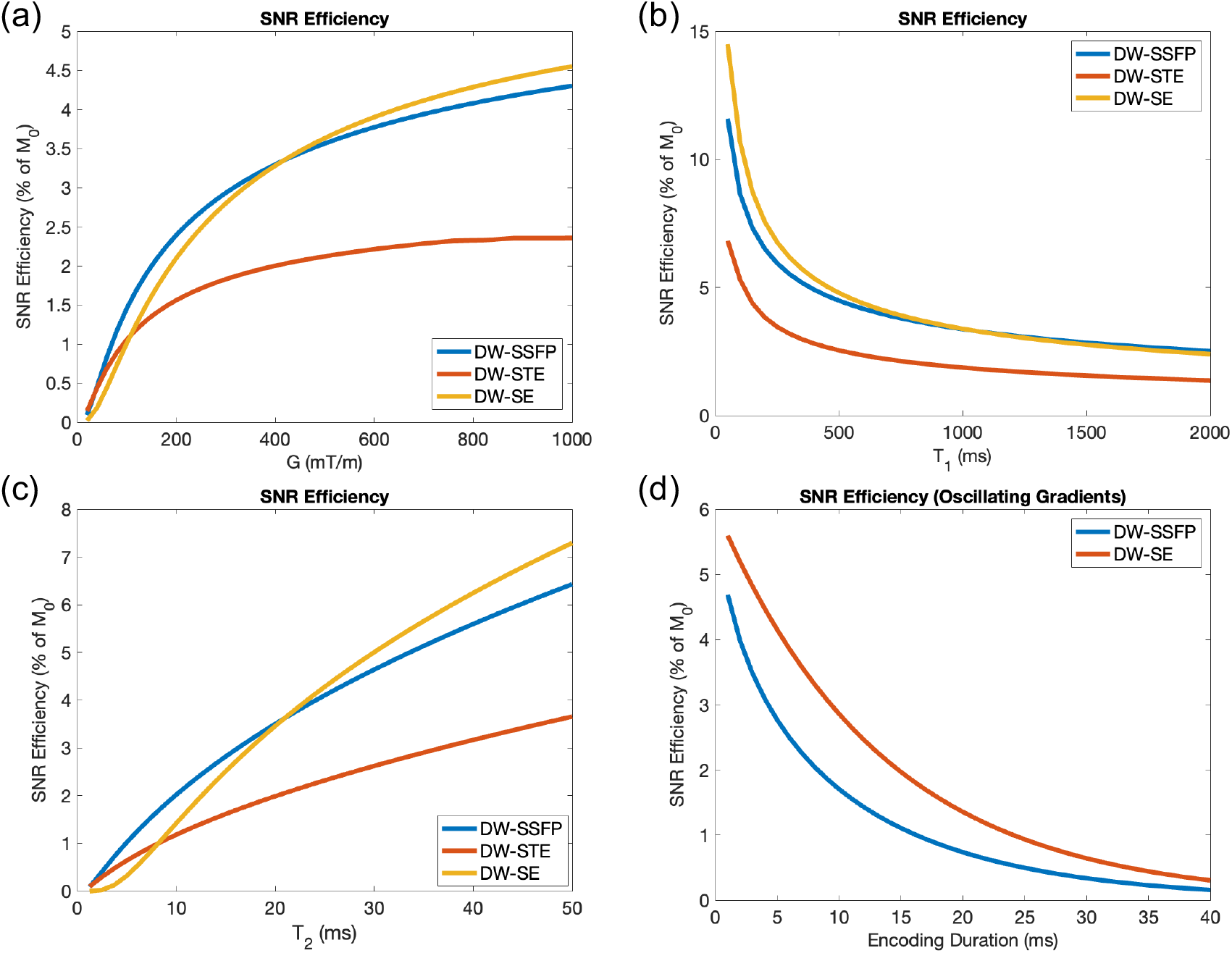
SNR-efficiency. SNR-efficiency estimation for DW-SSFP, DW-SE and DW-STE at a target b-value (*b* = 10 ms/μm^2^) as a function of gradient strength (a), T_1_ (b) and T_2_ (c). DW-SSFP predicts higher SNR-efficiency when compared to the DW-STE sequence across all investigated regimes, with the exception of very low gradient strengths. (a) The SNR-efficiency performance of DW-SSFP is reduced when compared to the DW-SE with increased gradient strength. (b) & (c) investigate the relationship between SNR-efficiency & relaxation regime at G = 1000 mT/m (rightmost point in (a)). DW-SSFP predicts higher SNR-efficiency with increased T_1_ and reduced T_2_ when compared to the DW-SE, motivating its use at ultra-high field. When considering oscillating gradients (d), DW-SSFP achieves reduced SNR-efficiency when compared to the DW-SE for matched encoding duration (per gradient). Simulations based on mean estimates of T_1_, T_2 &_ D from a cohort of post-mortem brains assuming free diffusion. Effective b-value for DW-SSFP based on matched levels of diffusion attenuation (corresponding to *b* = 10 ms/μm^2^ for DW-SE). For full information of implementation, see Methods. To explore how changing parameters influences these relationships, find the associated code [here](https://github.com/BenjaminTendler/SteadyStateDiffusionMicrostructure).

For oscillating gradients, DW-SSFP predicts reduced SNR-efficiency as a function of encoding duration when compared to the DW-SE in the investigated regime (Figure 10d). Whilst DW-SSFP can theoretically achieve higher diffusion-weighting with matched gradient waveforms (Figures 8b & d), these benefits could be mitigated by increasing the encoding period of the DW-SE sequence (e.g. having multiple repeats of a gradient waveform at a target frequency). These findings are broadly preserved when considering relaxation regimes found in post-mortem & in vivo tissue (Supporting Information Figure S8). The potential uses of DW-SSFP with oscillating gradients may be limited to investigational regimes where image distortions are particularly prominent.

## Discussion

### Interpreting DW-SSFP measurements

My framework shows that individual DW-SSFP measurements are equivalent to a distribution of signals summed across different diffusion regimes (Figures 5 and 6). This is in direct contrast to conventional diffusion MRI sequences that perform measurements at a single b-value or a single 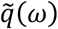 profile. Changes in DW-SSFP parameters modify the relative weighting of diffusion regimes that contribute to the measured signal (Figure 5 and Supporting Information Figure S5). This can be 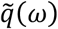 profiles, providing an experimental basis to characterise tissue microstructure.

Figures 5 and 6 additionally demonstrate that conventional DW-SSFP samples high b-value regimes and a narrow band of the diffusion spectrum compared to conventional diffusion imaging sequences. The time-*dependent* pathway framework demonstrates that DW-SSFP with oscillating diffusion gradients can probe different frequency components of the diffusion spectrum (Figure 7d), demonstrating a symmetric averaging of the power spectrum profile around the target frequency. Here, individual 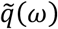 profiles are centred around the target frequency with small changes in their distribution shape, leading to a broad, Gaussian-like function.

The sequence parameters of DW-SSFP provide additional dimensions to probe tissue microstructure compared to conventional sequences that alter the diffusion regime through explicit changes to the diffusion-encoding module (i.e., q-value or diffusion time, Δ). For example, Figures 8a and b demonstrate that we can probe different diffusion regimes of a system by changing the acquisition flip angle, with no further changes to sequence timings. This effect can now be directly linked to modifications in the b-value distribution or power spectrum (Figures 5-7). The observation of flip-angle dependent diffusion weighting agrees with previous experimental work with DW-SSFP data acquired at multiple flip angles^[24,29]^ and EPG theory^[32]^.

Several attempts have been previously made to define a single effective b-value in steady-state diffusion imaging to facilitate comparisons with conventional diffusion sequences^[12,24,46]^. However, these b-value estimates have either (1) been derived based on estimated diffusion properties of the investigated system^[24]^ and/or (2) approximated the measured DW-SSFP signal^[12,14]^. The time-*independent* pathway framework shows that you cannot characterise the DW-SSFP signal using a single effective b-value (*b*_opp_), as there is no general solution for Eq. [10] where 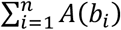⋅ Model(*b*_*i*_) = *A*_0_ ⋅ Model(*b*_eff_). However, by re-formulating in terms of a b-value distribution, we can characterise and interpret the measured signal with extremely high precision.

### Characterising tissue microstructure with DW-SSFP

My investigation finds that the signal-forming mechanisms of DW-SSFP predict conceptually higher levels of diffusion attenuation in small radii cylinders compared to a DW-SE sequence with identical gradient timings (Figures 8c). The SNR-efficiency properties of DW-SSFP (Figures 10a-c) reflects its potential for microstructural imaging when considering ultra-high field systems. Specifically, the theoretical SNR properties of DW-SSFP predict higher SNR-efficiency under matched diffusion attenuation when considering tissues with long T_1_ and short T_2_. These properties are reflected in the experimental dataset (Figure 9), where DW-SSFP achieves high SNR diffusivity estimates in the challenging imaging environment of fixed post-mortem tissue at 7T. Here the sequences high SNR-efficiency and strong diffusion-weighting address the low diffusivity^[47]^ and short-T_2_^[48]^ of fixed samples.

A key advantage DW-SSFP is that it acquires diffusion images with low or negligible distortions. Specifically, the short TR of DW-SSFP is compatible with a single-line readout, with no distortion correction applied to the experimental data in Figure 9 (three-line readout). At ultra-high field, increased distortions and short T_2_^*^ are particularly challenging when considering the EPI readouts associated with DW-SE. This suggests that DW-SSFP may still offer benefits for microstructural imaging in regimes where it achieves moderate reductions in SNR-efficiency when compared to the DW-SE.

When considering oscillating gradients, DW-SSFP is predicted to achieve a theoretical increase in diffusion attenuation (Figures 8b and d). This increase in diffusion attenuation is strongly dependent on the T_2_ and TR of the sample, where an increased T_2_ relative to the TR will lead to more high-signal pathways with that experience repeat sensitisation to gradient pairs. However, the relative SNR-efficiency of DW-SSFP is reduced when compared to the DW-SE (Figure 10d and Supporting Information Figure 8). Note that these findings do not extend to balanced steady-state sequences (e.g. TRUFI) with integrated oscillating gradients, which have not been investigated here.

The SNR-efficiency calculations presented here represent conceptual estimations of SNR-efficiency. They have not incorporated the impact of sequence dead time, readout duration limits, gradient duty cycle or T_2_^’^ decay. These properties are expected to vary depending on the precise implementation of the imaging sequence and properties of the MR system. In addition, the optimisations were based on free Gaussian diffusion, which does not consider the sequences specific sensitivity to microstructural features (e.g. restrictions). Understanding this potential warrants further investigation to perform quantitative comparisons of the sensitivity of DW-SSFP and DW-SE sequences individually optimised for microstructural sensitivity and SNR, and is the subject of ongoing work in my group^[49]^.

### Time-independent pathway framework

The time-*independent* framework demonstrates excellent agreement to analytical solutions and Monte Carlo simulations (Figures 8a and b). It provides high accuracy in comparison to existing approximation models of the DW-SSFP signal when T_2_ >> TR (Supporting Information Figure S4).

The time-*independent* framework exploits degeneracies in b-values to rapidly characterise the DW-SSFP signal. As the b-value distributions have relatively low dimensionality, we can pre-compute and store distributions to facilitate fitting of time-*independent* models to data. This approach was utilised for the tensor and NODDI^[39]^ estimation in Figure 9 (see Methods), demonstrating the utility of the framework as both a visualisation and parameter estimation tool. The experimental post-mortem comparison with diffusion tensor estimated parameters with excellent agreement to an analytical form (Figure 9a-c), demonstrating the accuracy of the proposed method.

A key feature of the time-*independent* pathway framework is that it can be extended to incorporate more sophisticated models based around compartments characterised by Gaussian diffusion without the requirement of extensive analytical derivations (Eqs. [10]-[13]). This is demonstrated via the integration of the NODDI model^[39]^ with experimental DW-SSFP data (Figure 9d), where no analytical form currently exists. However, the accuracy of the derived NODDI parameters is currently unclear. This challenge arises due to (1) the additional complexity of performing NODDI in fixed post-mortem tissue^[50]^, and (2) whether DW-SSFP measurements can be approximated as time-*independent* in the investigated experimental regimes. Specifically, as DW-SSFP probes a wide range of diffusion regimes simultaneously in a single measurement, whether the assumption of time-independence holds across different parameter spaces. This was not considered when acquiring the experimental DW-SSFP data used in this manuscript, which formed part of a larger, separate project^[28]^, and remain the subject of future investigation.

To the best of my knowledge, the analytical form of the DW-SSFP^[30]^ signal for (1) diffusion gradients of duration *δ* (Appendix 1), (2) oscillating gradients (Appendix 2) and (3) a tensor (Appendix 3) are presented here the first time in this format. Agreement with the proposed framework and Monte Carlo simulations demonstrate the robustness of these analytical forms.

### Time-dependent pathway framework

The proposed time-*dependent* framework provides a very general way to integrate existing models incorporating time-*dependent* effects that (1) does not require analytical solutions and (2) provides visualisation of the diffusion regimes being probed. These visualisations may support the identification of parameter spaces and novel gradient waveforms to probe distinct components of the diffusion spectrum (Figures 6 & 7).

The proposed time-*dependent* framework is based on the GPA^[41]^, which is typically considered valid in short and long diffusion time regimes. As a single DW-SSFP measurement probes multiple diffusion regimes simultaneously (Figures 5-7), a subset of pathways that contribute to the measured DW-SSFP signal may not be well approximated by the GPA. Despite this, results indicate that the time-*dependent* pathway framework gives good agreement to Monte Carlo simulations when investigating the parameter regimes and sample properties utilised in previous experimental DW-SSFP work^[29]^ (Figures 8c and d), with average errors of below 1%. Whilst I have not performed extensive investigations of the time-*dependent* pathway framework beyond the parameter regime explored in this article, the time-*dependent* pathway framework is not limited to the GPA, and could be readily extended to more advanced models of the diffusion MRI signal^[51]^.

A key limitation of the time-*dependent* pathway framework is that each signal-forming pathway needs to be characterised separately to estimate diffusion attenuation, leading to long computational times. Whilst this problem can be parallelised, the number of pathways that contribute to a given measurement are extremely high (Supporting Information Figure S6), limiting the space of investigation. Here I address this by limiting the investigation to pathways that experience up to four TRs in the transverse plane. This accounts for ∼99% of the measured DW-SSFP signal for the parameters investigated here (Figure S6). However, this approximation would lead to amplified errors for investigations of tissue with long T_2_ or acquisitions with short TRs (Supporting Information Figure S4), and limits the framework to estimating & visualising the DW-SSFP signal, rather than its utility for parametric fits (model inversion). An approach to identify the most meaningful pathways that contribute to a measurement would facilitate extending the time-*dependent* framework to higher order pathways, but no robust method to achieve this was identified as part of this work. A key challenge is that the signal contributed from pathways that experience several TRs in the transverse plane is the average of many pathways with positive and negative signal amplitudes (Supporting Information Figure S9).

An alternative approach to investigate time-*dependent* models is to utilise Monte Carlo simulations, which can be used to investigate the integration of sophisticated microstructural systems with the DW-SSFP sequence (Figure 7b). Taken together, Monte Carlo simulations may provide the most generalisable approach to characterise time-*dependent* systems with DW-SSFP across a large parameter space, despite the long simulation times required to model the approach of the DW-SSFP signal to a steady-state (typically requiring 10s to 100s of TRs – Supporting Information Figure S10). The proposed time-*dependent* pathway framework would provide complementary information to visualise the diffusion regimes investigated, in addition to gaining both understanding and intuition of the signal-forming mechanisms of DW-SSFP.

## Conclusion

I established a framework providing forward predictions of the measured DW-SSFP signal under a broad range of existing biophysical models incorporating time-*independent* and -*dependent* diffusion phenomena. The framework facilitates the visualisation and characterisation of the measured DW-SSFP signal, providing both understanding and intuition of how sequence properties influence diffusion attenuation. When considering time-*independent* diffusion, the framework additionally provides an approach to integrate and estimate model parameters from experimental data, without the requirement of model-specific derivations. Findings demonstrate excellent agreement with existing analytical and simulation methods, demonstrating that DW-SSFP achieves microstructural sensitivity by probing multiple diffusion regimes simultaneously. Combined with its theoretical SNR-efficiency and low-distortion properties, the DW-SSFP sequence may offer considerable potential for characterising tissue microstructure, particularly at ultra-high field.

## Methods

### Default parameters and software

Default sequence parameters for all simulations (unless explicitly stated) were matched to the experimental DW-SSFP analysis at 7T performed in Tendler et al.^[29]^, setting G = 52 mT/m, *δ* = 13.56 ms, *α* = 24° and TR = 28 ms, with *D* = 0.2 μm^2^/s, T_1_ = 600 ms and T_2_ = 40 ms. Investigations were performed using MATLAB (2023a, The MathWorks, Inc., Natick, MA) on a Macbook Pro (macOS Big Sur, M1 chip, 16GB ram).

For the oscillating gradient investigations, parameters were matched to Aggarwal et al.^[45]^, setting *G* = 674 mT/m, *δ* = 20 ms and TR/Δ = 25 ms. All other parameters were matched to those defined in the previous paragraph.

### Time-independent framework

Details of the time-*independent* framework are provided in the Theory section. For the practical implementation, a pathway was characterised as contributing to the measured signal if it had an amplitude (fraction of M_0_) above a defined threshold, here set to 10^-28^. I excluded pathways that persisted for over 1000 TRs (maximum relaxation attenuation O(10^−24^)) and dictionary entries persisting for more than 10 TRs in the transverse plane (maximum relaxation attenuation O(10^−4^)). These thresholds were found to have a negligible impact on signal estimation in the investigated regimes. Gradient waveforms were discretised into 1000 time points per TR.

### Time-dependent framework

Details of the time-*dependent* framework are provided in the Theory section. For the practical implementation, I modelled diffusion attenuation using the GPA as described in Eq. [14]. This was implemented using the approach described in Ning et al.^[42]^, reparameterising the GPA as a function of the mean-squared displacement of the diffusion system and the autocorrelation function of the gradient waveform. This approach was chosen as it can more rapidly estimate the signal attenuation when compared to Eq. [14]. An analytical form of the mean squared displacement of a cylinder was defined based on Equation 32 in Mortensen et al^[52]^.

I characterised a pathway as contributing to the measured signal if it had an amplitude (fraction of M_0_) above a defined threshold, here set to 10^-10^. I excluded pathways that persisted for over 1000 TRs (maximum relaxation attenuation O(10^−24^)) or more than 4 TRs (maximum relaxation attenuation O(10^−2^)) in the transverse plane, corresponding to ∼99% of the measured signal for the investigated parameters (Supporting Information Figure S6). Gradient waveforms were discretised into 100 time points per TR. The average time to estimate a single data point for the cylindrical restriction investigation (Figures 8c and d) was approximately 70 seconds.

### Monte Carlo simulations

Spin trajectories for all Monte Carlo simulations were generated using the Camino *datasynth* function^[53]^, modified to produce trajectories that followed a Gaussian distribution of displacements per time-step. The DW-SSFP & DW-SE signal was subsequently modelled using custom MATLAB code (2023a, The MathWorks, Inc., Natick, MA). All simulations used 5 ⋅ 10^5^ spins.

For DW-SSFP, I simulated 10,000 time-steps (100 time-steps per TR, 100 TRs). These parameters ensured the measured signal reached a steady state (Supporting Information Figure S10). for DW-SE, I simulated 200 time-steps (200 time-steps per TR, 1 TR). For the DW-SSFP simulations, the initial position of spins was modified to match the 2π dephasing condition of DW-SSFP.

### Experimental DW-SSFP acquisition and analysis

Acquisition of the experimental data used in this manuscript is described in Tendler et al^[29]^. Briefly, DW-SSFP data of a whole, human post-mortem brain was acquired with the following parameters: resolution = 850 μm iso., no. directions = 120, q = 300 cm^−1^, G = 52 mT/m, *δ* = 13.56 ms, *α* = 24°& 94°, TE = 21 ms, TR = 28 ms, Bandwidth = 393 Hz/mm, Acquisition time per direction = 5m 47s, acquisition time per flip angle = ∼ 12h. Voxelwise quantitative T_1_, T_2_ and B_1_ maps were additionally derived using complementary acquisitions due to DW-SSFP’s dependence on relaxation times and flip angle (Figure 2). The original processed data associated with this project can be accessed via the Digital Brain Bank^[54]^ (*Human ALS MRI-Histology dataset*).

For fitting to experimental data, I used the time-*independent* framework to generate a b-value distribution dictionary for a range of T_1_ (100 to 1000 ms; 10 ms increments), T_2_ (1 to 60 ms; 1 ms increments), and flip angles (1° to 140°; 1° increments). The final dictionary contained 764,400 b-value distributions, requiring ∼20 GB of space. This dictionary was generated over several days on a Macbook Pro (macOS Big Sur, M1 chip, 16GB ram). However, this process can be readily parallelised and only need to be performed once for a given set of DW-SSFP parameters.

Fitting for the time-*independent* pathway framework was performed using the b-value distribution dictionary and Eq. [10] integrating the tensor model (Eq. [12]). The comparison analytical DW-SSFP model^[30]^ with integrated tensor is described in Appendix 3. Fitting was performed with MATLAB (2023a, The MathWorks, Inc., Natick, MA) using *lsqnonlin* on my centre’s computing cluster.

For the experimental NODDI investigation, I similarly integrated the b-value distribution dictionary with Eq. [10] incorporating the NODDI model^[39]^ (Eq. [13]). Parameter estimation was performed via a multi-step pipeline, reflecting the implementation steps performed by the [NODDI toolbox](https://www.nitrc.org/projects/noddi_toolbox/). Briefly, the principal fibre orientation 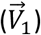 was first initialised based on a tensor 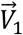 estimate. The isotropic volume fraction (*f*_*iso*_), intracellular volume fraction (*f*_*intra*_) and orientation dispersion (*OD*) were subsequently estimated based on a custom grid-search fitting algorithm. To account for fixed post-mortem tissue, the intrinsic free diffusion coefficient (*D*_||_) was fixed to 0.42 μm^2^/ms, estimated based on a custom implementation of the spherical mean technique^[55]^ for DW-SSFP data. The isotropic diffusion coefficient (*D*_*iso*_) was fixed to 2 μm^2^/ms. The remaining free parameters (*f*_*iso*_, *f*_*intra*_, *OD* and 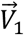 were simultaneously estimated using MATLAB *lsqnonlin*. Fitting was performed using a combination of custom MATLAB code and functions from the NODDI toolbox. It was not possible to investigate the existence of a dot compartment based on the acquired DW-SSFP data, with incorporation leading to noisy parameter estimates.

### Ethics statement

Experimental data derived from a whole, human post-mortem brain (Figure 9) used tissue provided by the Oxford Brain Bank, a research ethics committee (REC) approved, HTA regulated research tissue bank. The study was conducted under the Oxford Brain Bank’s generic Research Ethics Committee approval (15/SC/0639).

### SNR-Efficiency Comparison

SNR-efficiency was defined as:

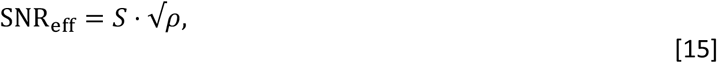

Where *S* is the estimated signal amplitude from the DW-SSFP, DW-SE and DW-STE sequence (Appendix 4), and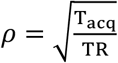, where T_acq_ is the readout duration of each sequence (Appendix 5).

Estimation of parameters that maximised SNR_eff_ at a target b-value was performed using the *lsqnonlin* function in MATLAB (2023a, The MathWorks, Inc., Natick, MA), assuming free Gaussian diffusion. For conventional DW-SSFP (Figures 10a-c), optimisation was performed for a target b-value by optimising:

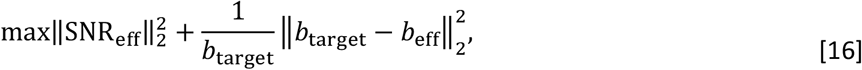

where *b*_target_ was fixed to 10 ms/μm^2^ and *b*_eff_ is an effective DW-SSFP b-value that achieves the same level of diffusion attenuation as a conventional DW-SE or DW-STE sequence:

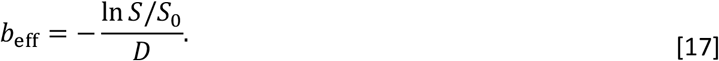

Optimisations were performed assuming no dead time, readout duration limits, gradient duty cycle limits or T_2_^’^ decay. These properties are expected to vary depending on the precise implementation of the imaging sequence and properties of the MR system.

For the optimisation, T_1_, T_2 &_ D were set equal to 552 ms, 26.8 ms and 0.14 μm^2^/ms, the median values estimated in white matter at 7T from a cohort of post-mortem brains fixed with neutral buffered formalin^[56]^. This cohort included the brain used for the Tensor and NODDI investigation (Figure 9).

## Appendices

### Appendix 1

Based on the proposed analytical model in Freed et al.^[30]^, the DW-SSFP sequence incorporating a diffusion gradient of duration *δ* is modelled as:

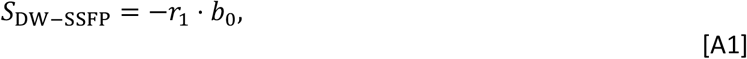

where:

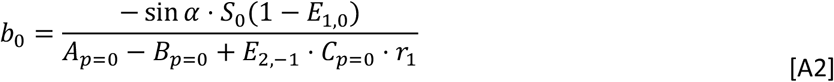

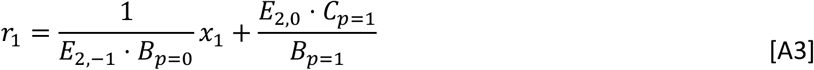

defining:

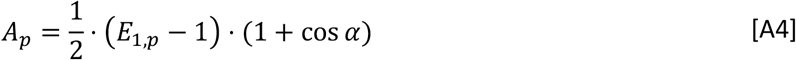

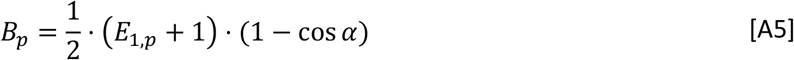

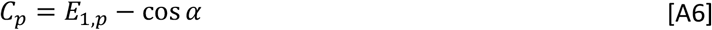

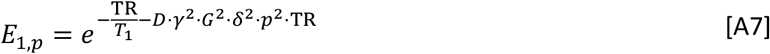

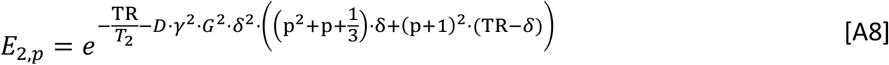

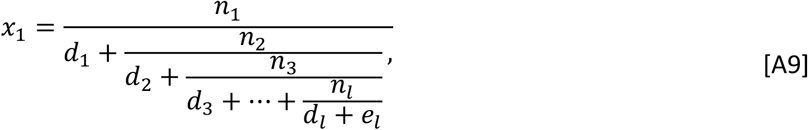

with:

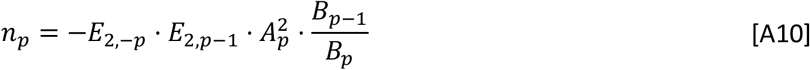

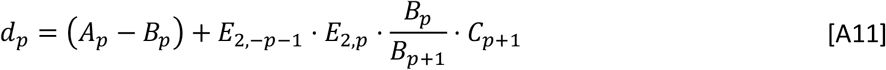

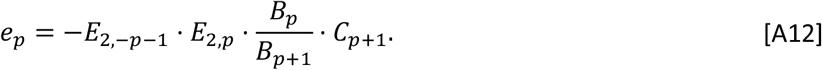

Find the associated code for this model [here](https://github.com/BenjaminTendler/SteadyStateDiffusionMicrostructure).

### Appendix 2

The DW-SSFP Analytical model for oscillating gradients is equivalent to Appendix 1, setting:

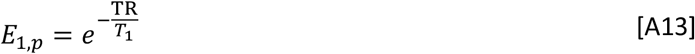

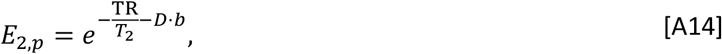

where *b* is equivalent to a b-value of a single instance of the oscillating gradient waveform (Figure 7a; here *b* = 0.35 ms/μm^2^). Find the associated code for this model [here](https://github.com/BenjaminTendler/SteadyStateDiffusionMicrostructure).

### Appendix 3

The DW-SSFP Analytical model with integrated tensor is equivalent to Appendix 1, setting:

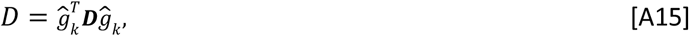

where ĝ_*k*_is the gradient orientation and ***D*** is the diffusion tensor. Find the associated code for this model [here](https://github.com/BenjaminTendler/SteadyStateDiffusionMicrostructure).

### Appendix 4

*S*_*D*W −SSFP_is defined in Appendix 1, with

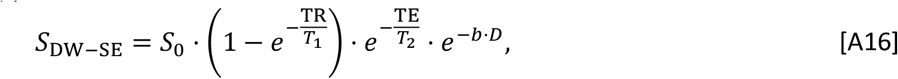

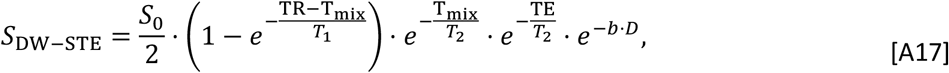

where T_mix_ is the mixing time.

### Appendix 5

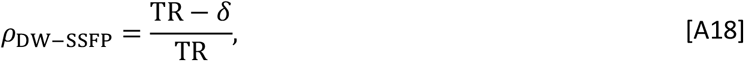

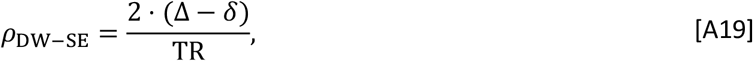

where Δ is the diffusion time, defining TE = 2 ⋅ Δ (i.e. no dead time between the RF pulse and the start of the second diffusion gradient), and

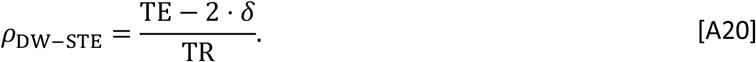

## Additional Information

BCT declares no competing interests.

## Acknowledgements

BCT would like to thank Karla L. Miller and Shaihan J. Malik for helpful discussions during the development of this project and manuscript feedback.

BCT is funded by a Sir Henry Wellcome Postdoctoral Fellowship (Wellcome Trust) [222829/Z/21/Z]. The Wellcome Centre for Integrative Neuroimaging is supported by core funding from the Wellcome Trust [203139/Z/16/Z and 203139/A/16/Z]. Human tissue was provided by the Oxford Brain Bank, funded by the Medical Research Council, Brains for Dementia Research, and the NIHR Oxford Biomedical Research Centre.

This research was funded in whole, or in part, by the Wellcome Trust [222829/Z/21/Z]. For the purpose of open access, the author has applied a CC BY public copyright licence to any Author Accepted Manuscript version arising from this submission.

## Data Availability

Software for the time-*independent* and -*dependent* frameworks, including scripts to replicate many of the findings presented in this manuscript, are available here. The original processed post-mortem data associated with this project can be accessed via the Digital Brain Bank^[54]^ (*Human ALS MRI-Histology dataset*).

## Author Contribution Statement

BCT was responsible for the described investigation.

