## Supporting Information for "Investigating tissue microstructure using steady-state diffusion MRI"

### Characterising the DW-SSFP signal: Examples

*Example 1: Identifying signal-forming pathways*

Prior to excitation, all magnetisation is longitudinal ($\tilde{Z}_0$). In Eq. [1] (Main Text), magnetisation ($\boldsymbol{M}$) is initialised in the transverse plane at the end of the first TR, defining:

$$\boldsymbol{M}_{\mathbf{init}} = \begin{bmatrix} -i \cdot E_2 \cdot e^{i\phi} \cdot \sin\alpha \\ i \cdot E_2 \cdot e^{-i\phi} \cdot \sin\alpha \\ 0 \end{bmatrix}; \ \boldsymbol{F}_{\mathbf{init}} = \begin{bmatrix} \tilde{F}_{+1} \\ \tilde{F}_{-1} \\ \tilde{Z}_0 \end{bmatrix}, \qquad \text{[S1]}$$

After a single application of the $\boldsymbol{ET} \circledast$ operator and diffusion gradient:

$$\boldsymbol{ET} \circledast \boldsymbol{M}_{\mathbf{init}} = \begin{bmatrix} -E_2^2 \cdot i \cdot e^{i\phi} \cdot \cos^2\frac{\alpha}{2} \cdot \sin\alpha & E_2^2 \cdot i \cdot e^{i\phi} \cdot \sin^2\frac{\alpha}{2} \cdot \sin\alpha & 0 \\ -E_2^2 \cdot i \cdot e^{-i\phi} \cdot \sin^2\frac{\alpha}{2} \cdot \sin\alpha & E_2^2 \cdot i \cdot e^{-i\phi} \cdot \cos^2\frac{\alpha}{2} \cdot \sin\alpha & 0 \\ -E_1 \cdot E_2 \cdot \frac{1}{2} \cdot \sin^2\alpha & -E_1 \cdot E_2 \cdot \frac{1}{2} \cdot \sin^2\alpha & 0 \end{bmatrix}, \qquad \text{[S2]}$$

corresponding to the following states:

$$(\boldsymbol{ET} \circledast \boldsymbol{M}_{\mathbf{init}})_{\text{States}} = \begin{bmatrix} \tilde{F}_{+2} & \tilde{F}_0^* & \tilde{F}_{+1} \\ \tilde{F}_0 & \tilde{F}_{-2} & \tilde{F}_{-1} \\ \tilde{Z}_{+1} & \tilde{Z}_{-1} & \tilde{Z}_0 \end{bmatrix}. \qquad \text{[S3]}$$

$\tilde{F}_0$ and $\tilde{F}_0^*$ states are complex-conjugates, and I define the $\tilde{F}_0$ state as contributing to the measured signal. Here, we identify a single $\tilde{F}_0$ state with signal amplitude $= -E_2^2 \cdot i \cdot e^{-i\phi} \cdot \sin^2\frac{\alpha}{2} \cdot \sin\alpha$. This corresponds to a signal-forming 'spin-echo' pathway evolving via $\tilde{Z}_0 \rightarrow \tilde{F}_{+1} \rightarrow \tilde{F}_0$.

*Example 2: Identifying further signal-forming pathways*

After one application of the $\boldsymbol{ET} \circledast$ operator, we identified a single $\tilde{F}_0$ state (Equation S3). We can identify additional signal-forming pathways with repeat application of the $\boldsymbol{ET} \circledast$ operator.

Building on Equations S2 & S3:

$$\boldsymbol{ET} \circledast \boldsymbol{ET} \circledast \boldsymbol{M}_{\mathbf{init}} =$$

$$\begin{bmatrix} -E_2^3 \cdot i \cdot e^{i\phi} \cdot \cos^4\frac{\alpha}{2} \cdot \sin\alpha & E_2^3 \cdot i \cdot e^{i\phi} \cdot \cos^2\frac{\alpha}{2} \cdot \sin\alpha \cdot \sin^2\frac{\alpha}{2} & 0 & -E_2^3 \cdot i \cdot e^{i\phi} \cdot \sin^4\frac{\alpha}{2} \cdot \sin\alpha & E_2^3 \cdot i \cdot e^{i\phi} \cdot \cos^2\frac{\alpha}{2} \cdot \sin\alpha \cdot \sin^2\frac{\alpha}{2} & 0 & E_1 \cdot E_2^2 \cdot \frac{i}{2} \cdot e^{i\phi} \cdot \sin^3\alpha & E_1 \cdot E_2^2 \cdot \frac{i}{2} \cdot e^{i\phi} \cdot \sin^3\alpha & 0 \\ -E_2^3 \cdot i \cdot e^{-i\phi} \cdot \cos^2\frac{\alpha}{2} \cdot \sin\alpha \cdot \sin^2\frac{\alpha}{2} & E_2^3 \cdot i \cdot e^{-i\phi} \cdot \sin^4\frac{\alpha}{2} \cdot \sin\alpha & 0 & -E_2^3 \cdot i \cdot e^{-i\phi} \cdot \cos^2\frac{\alpha}{2} \cdot \sin\alpha \cdot \sin^2\frac{\alpha}{2} & E_2^3 \cdot i \cdot e^{-i\phi} \cdot \cos^4\frac{\alpha}{2} \cdot \sin\alpha & 0 & -E_1 \cdot E_2^2 \cdot \frac{i}{2} \cdot \square\square^{-i\phi} \cdot \sin^3\alpha & -E_1 \cdot E_2^2 \cdot \frac{i}{2} \cdot e^{-i\phi} \cdot \sin^3\alpha & 0 \\ -E_1 \cdot E_2^2 \cdot \frac{1}{2} \cdot \cos^2\frac{\alpha}{2} \cdot \sin^2\alpha & E_1 \cdot E_2^2 \cdot \frac{1}{2} \cdot \sin^2\alpha \cdot \sin^2\frac{\alpha}{2} & 0 & E_1 \cdot E_2^2 \cdot \frac{1}{2} \cdot \sin^2\alpha \cdot \sin^2\frac{\alpha}{2} & -E_1 \cdot E_2^2 \cdot \frac{1}{2} \cdot \cos^2\frac{\alpha}{2} \cdot \sin^2\alpha & 0 & -E_1^2 \cdot E_2 \cdot \frac{1}{2} \cdot \cos\alpha \cdot \sin^2\alpha & -E_1^2 \cdot E_2 \cdot \frac{1}{2} \cdot \cos\alpha \cdot \sin^2\alpha & 0 \end{bmatrix}, \qquad \text{[S4]}$$

$$(\boldsymbol{ET} \circledast \boldsymbol{ET} \circledast \boldsymbol{M}_{\mathbf{init}})_{\mathbf{States}} = \begin{bmatrix} \tilde{F}_{+3} & \tilde{F}_{+1} & \tilde{F}_{+2} & \tilde{F}_{+1} & \tilde{F}_{-1} & \tilde{F}_0^* & \tilde{F}_{+2} & \tilde{F}_0^* & \tilde{F}_{+1} \\ \tilde{F}_{+1} & \tilde{F}_{-1} & \tilde{F}_0 & \tilde{F}_{-1} & \tilde{F}_{-3} & \tilde{F}_{-2} & \tilde{F}_0 & \tilde{F}_{-2} & \tilde{F}_{-1} \\ \tilde{Z}_{+2} & \tilde{Z}_0 & \tilde{Z}_{+1} & \tilde{Z}_0 & \tilde{Z}_{-2} & \tilde{Z}_{-1} & \tilde{Z}_{+1} & \tilde{Z}_{-1} & \tilde{Z}_0 \end{bmatrix} \qquad \text{[S5]}$$

After two applications of the $\boldsymbol{ET} \circledast$ operator, we identify two $\tilde{F}_0$ states with signal amplitudes $= 0$ & $-E_1 \cdot E_2^2 \cdot \frac{i}{2} \cdot e^{-i\phi} \cdot \sin^3\alpha$. These correspond to a signal-forming 'delayed spin-echo' pathway evolving via $\tilde{Z}_0 \rightarrow \tilde{Z}_0 \rightarrow \tilde{F}_{+1} \rightarrow \tilde{F}_0$ and a stimulated echo pathway evolving via $\tilde{Z}_0 \rightarrow \tilde{F}_{+1} \rightarrow \tilde{Z}_{+1} \rightarrow \tilde{F}_0$. As noted in the Theory (Main Text), delayed pathways are accounted for within the initialisation term (Eq. [1] - Main Text) and therefore have signal amplitude = 0.

We can recursively apply the $\boldsymbol{ET} \circledast$ operator to identify additional signal-forming pathways.

### Supporting Information Table

| Figure | $n_{long}$ | $A$ | $b$ |
|---|---|---|---|
| (a) | 1 | $A_{\text{init}} \cdot (E_1 \cdot \cos\alpha)^{(n-1)}$ | $\gamma^2 G^2 \delta^2 \cdot k_n^2 \cdot (n-1) \cdot \text{TR} + b_{\text{init}}$ |
| (b) | 2 | $A_{\text{init}} \cdot (E_1 \cdot \cos\alpha)^{(n+m-2)}$ | $\gamma^2 G^2 \delta^2 \cdot (k_n^2 \cdot (n-1) + k_m^2 \cdot (m-1)) \cdot \text{TR} + b_{\text{init}}$ |
| (c) | 3 | $A_{\text{init}} \cdot (E_1 \cdot \cos\alpha)^{(n+m+o-3)}$ | $\gamma^2 G^2 \delta^2 \cdot (k_n^2 \cdot (n-1) + k_m^2 \cdot (m-1) + k_o^2 \cdot (o-1)) \cdot \text{TR} + b_{\text{init}}$ |

| Figure | $A_{\text{init}}$ | $b_{\text{init}}$ | $k_{n,m,o\ldots}$ |
|---|---|---|---|
| (a) | $-E_1 \cdot E_2^2 \cdot \frac{i}{2} \cdot e^{-i\phi} \cdot \sin^3\alpha$ | $\gamma^2 G^2 \delta^2 \cdot \left(2 \cdot \text{TR} - \frac{\delta}{3}\right)$ | $k_n = 1$ |
| (b) | $-E_1^2 \cdot E_2^6 \cdot \frac{i}{4} \cdot e^{-i\phi} \cdot \sin^5\alpha \cdot \cos^4\frac{\alpha}{2} \cdot \sin^2\frac{\alpha}{2}$ | $\gamma^2 G^2 \delta^2 \cdot (19 \cdot \text{TR} - \delta)$ | $k_n = 2, k_m = 2$ |
| (c) | $-E_1^3 \cdot E_2^{10} \cdot \frac{i}{8} \cdot e^{-i\phi} \cdot \sin^7\alpha \cdot \cos^{12}\frac{\alpha}{2}$ | $\gamma^2 G^2 \delta^2 \cdot (135 \cdot \text{TR} - \frac{5 \cdot \delta}{3})$ | $k_n = 5, k_m = 4, k_o = 3$ |

Table S1: **Properties of the three example dictionary components displayed in Figure 4 (Main Text)**. $A_{\text{init}}$ estimated using the ***ET*** operator (Eq. [2]; Main Text). $b_{\text{init}}$ derived using the approach in Weigel et al.[1]. Here, $n_{long}$ = number of distinct longitudinal periods. $n, m, o$ defined in Figure 4 (Main Text).

### Supporting Information Figures

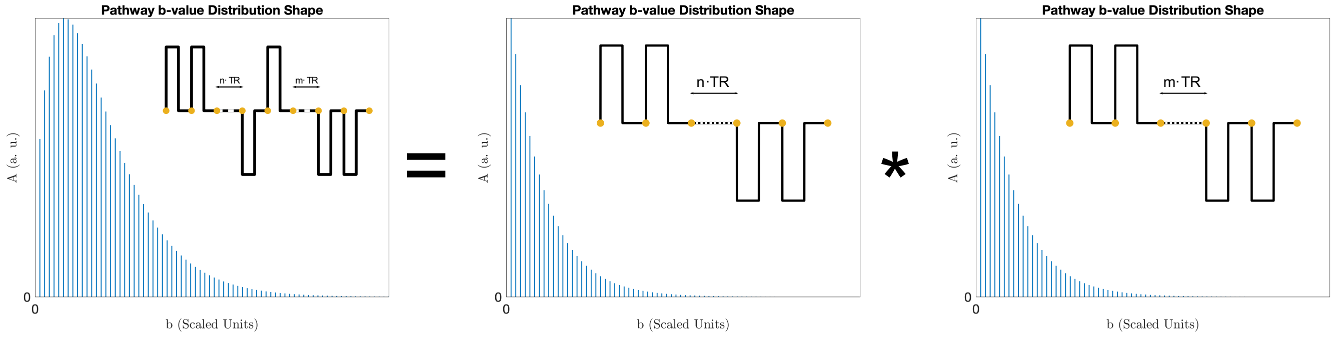


Figure S1: **b-value distribution shape estimation.** The b-value distribution of dictionary entries with two longitudinal periods ($n_{long} = 2$) are mathematically linked to dictionary entries with one longitudinal period ($n_{long} = 1$) via a convolution (Eq. [7] – Main Text). This property can be extended to dictionary entries with several longitudinal periods (Eq. [8] – Main Text), providing a rapid approach to synthesise b-value distributions. Left: b-value distribution with $n_{long} = 2$, $k_n = 2$ & $k_m = 2$. Right: Two identical b-value distributions with $n_{long} = 1$ and $k_n = 1$. Synthesised b-value distribution shape equivalent to dictionary entry in Figure 4b, where Figure 4b has been additionally shifted along the x-axis by $b_{\text{init}}$.

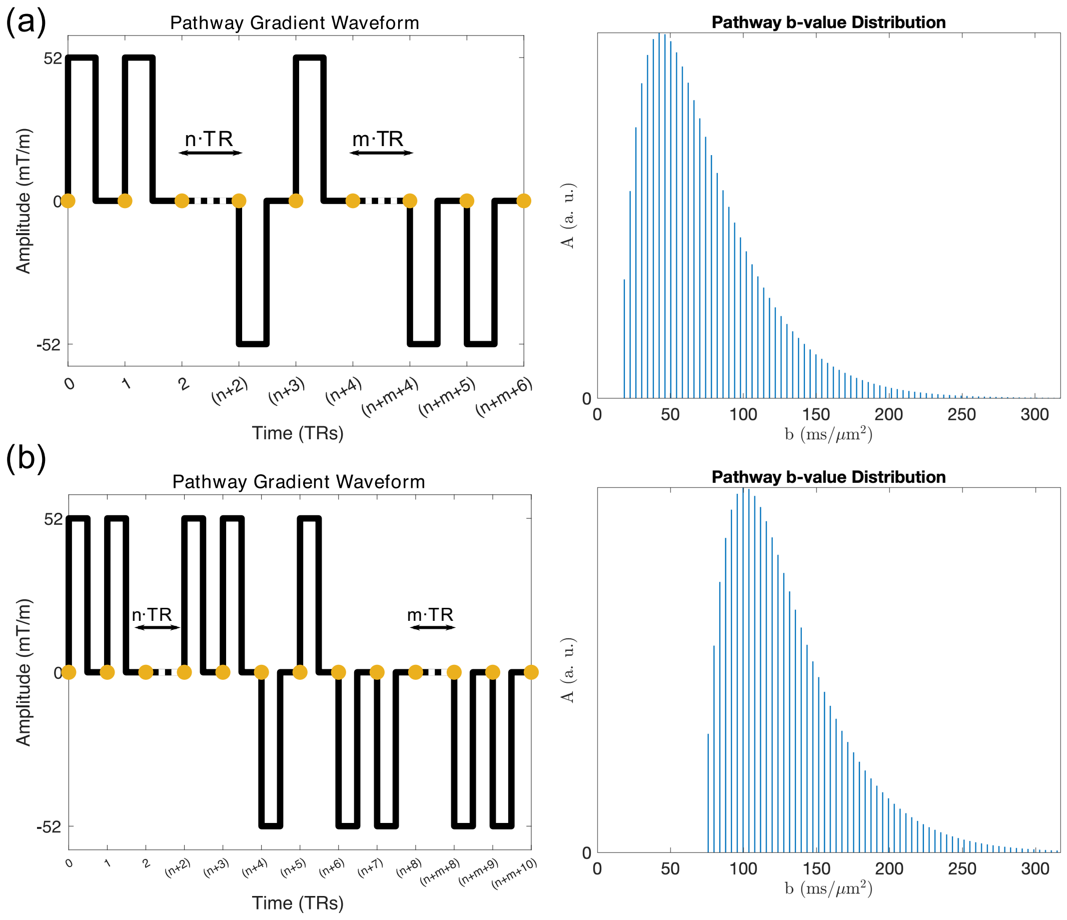


Figure S2: **b-value distribution degeneracy.** The b-value distribution shape for a given gradient waveform is associated with the (1) number ($n_{long}$) and (2) phase state ($k_{l,m,\ldots}$) of longitudinal periods. Whilst the gradient waveforms in (a) and (b) appear distinct, they both have two longitudinal periods ($n_{long} = 2$) associated with phase states $k_n = k_m = 2$. Subsequent b-value distribution shapes are identical, initialised along the x-axis by the b-value of the shortest gradient waveform $b_{\text{init}}$ ($\propto (19 \cdot \mathrm{TR} - \delta)$ and $(77 \cdot \mathrm{TR} - 5 \cdot \frac{\delta}{3})$ for (a) and (b)). Rather than calculating the b-value distribution of each dictionary entry individually, we only need to identify the unique b-value shape associated with different combinations of $n_{long}$ and $k_{n,m,o,\ldots}$.

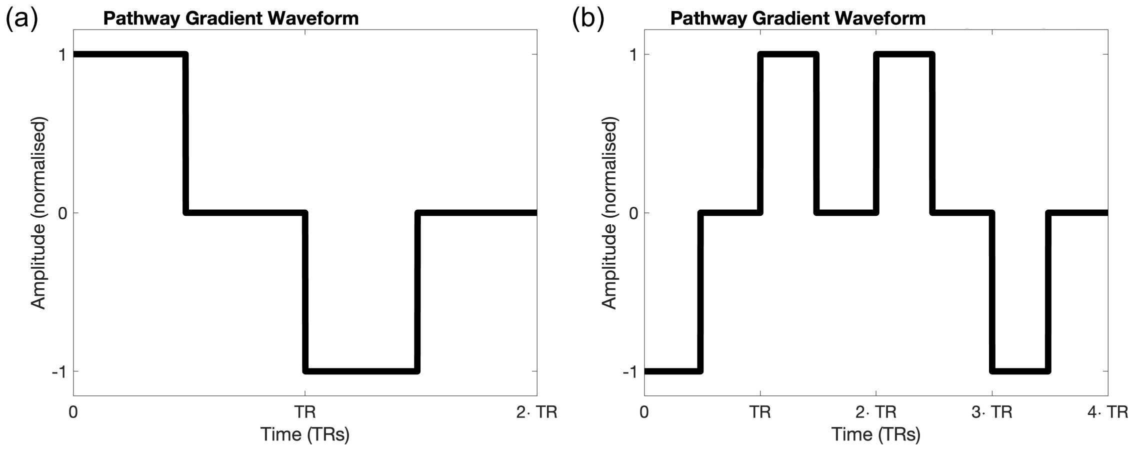


Figure S3: **Signal-forming pathways with positive & negative signal amplitudes**. Example gradient waveforms experienced by two signal-forming pathways that give rise to signal amplitudes with opposite signs. (a) corresponds to a spin-echo pathway evolving via $\tilde{Z}_0 \to \tilde{F}_{+1} \to \tilde{F}_0$, with a corresponding signal amplitude of $-i \cdot E_2^2 \cdot e^{-i\phi} \cdot \sin^2\frac{\alpha}{2} \cdot \sin\alpha$. (b) corresponds to a dual spin-echo pathway evolving via $\tilde{Z}_0 \to \tilde{F}_{-1} \to \tilde{F}_0^* \to \tilde{F}_{+1} \to \tilde{F}_0$, with a corresponding signal amplitude of $i \cdot E_2^4 \cdot e^{-i\phi} \cdot \sin^4\frac{\alpha}{2} \cdot \cos^2\frac{\alpha}{2} \cdot \sin\alpha$.

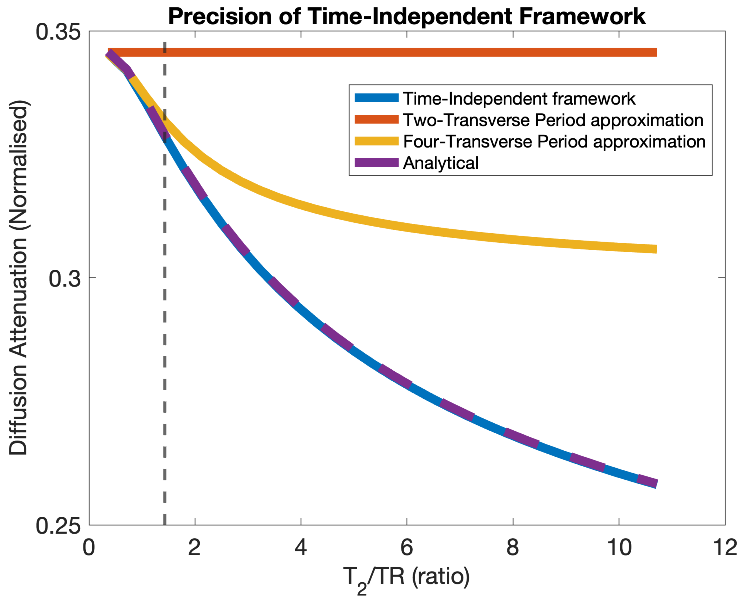


Figure S4: **Comparison of the time-*independent* framework with analytical and approximation DW-SSFP models**. The time-*independent* framework (blue) gives excellent agreement to an analytical DW-SSFP model (purple) (Eq. A1 – Main Text) across a range of $T_2$/TR ratios. The two-transverse period approximation (red) is independent of $T_2$, rapidly deviating from the analytical model (purple). The four-transverse period approximation (yellow) gives good agreement at low $T_2$/TR ratios, with increasing deviation at higher values. The dashed vertical line indicates the $T_2$/TR ratio used for the investigation in this manuscript., with parameters based on post-mortem DW-SSFP investigations at 7T as described in Tendler et al.[2] (defined in Methods). Here, the x-axis spans $T_2$ values from 10 to 300 ms. To explore how changing parameters impacts the precision of different methods, find the associated code here.

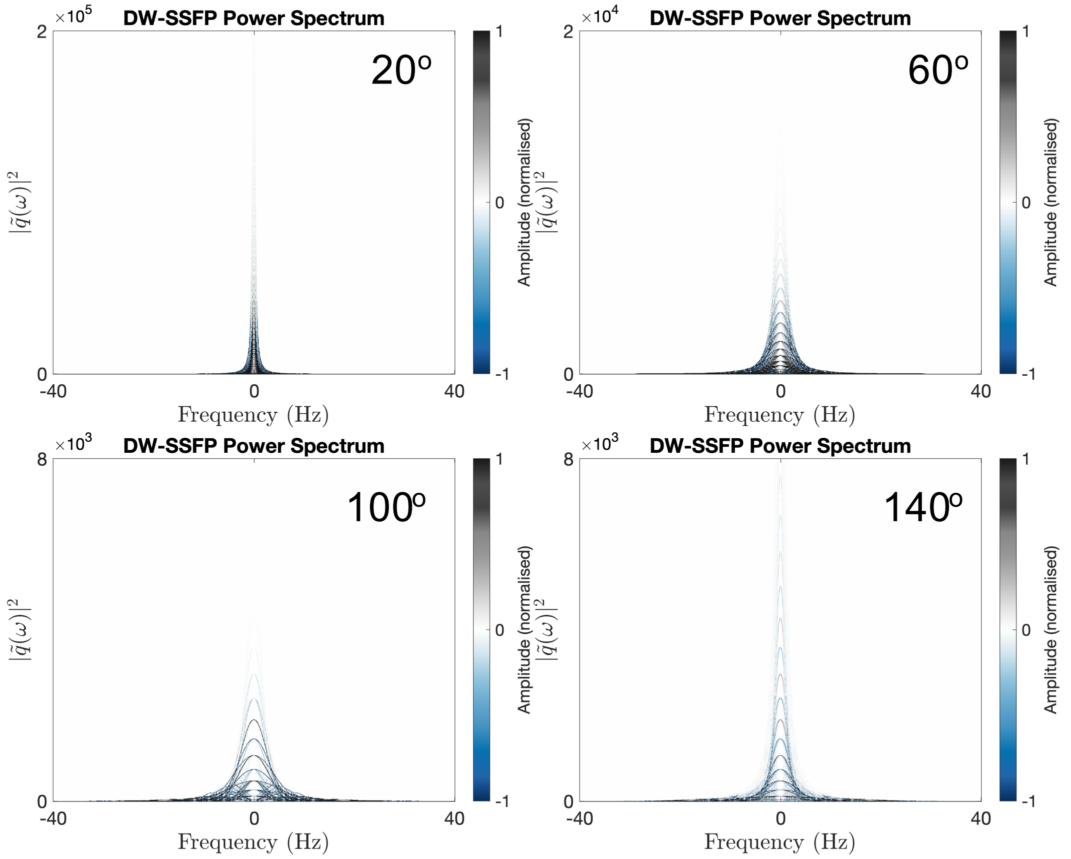


Figure S5: **Change in power spectrum density weighting of the DW-SSFP signal with flip angle**. Complementary to Figure 6 (main text), here we observe that an increase in flip angle is associated with a change in amplitude and width of the dominant power spectrum components. Sequence parameters based on post-mortem DW-SSFP investigations at 7T as described in Tendler et al.[2] (defined in Methods). Note that the y axes span different extents.

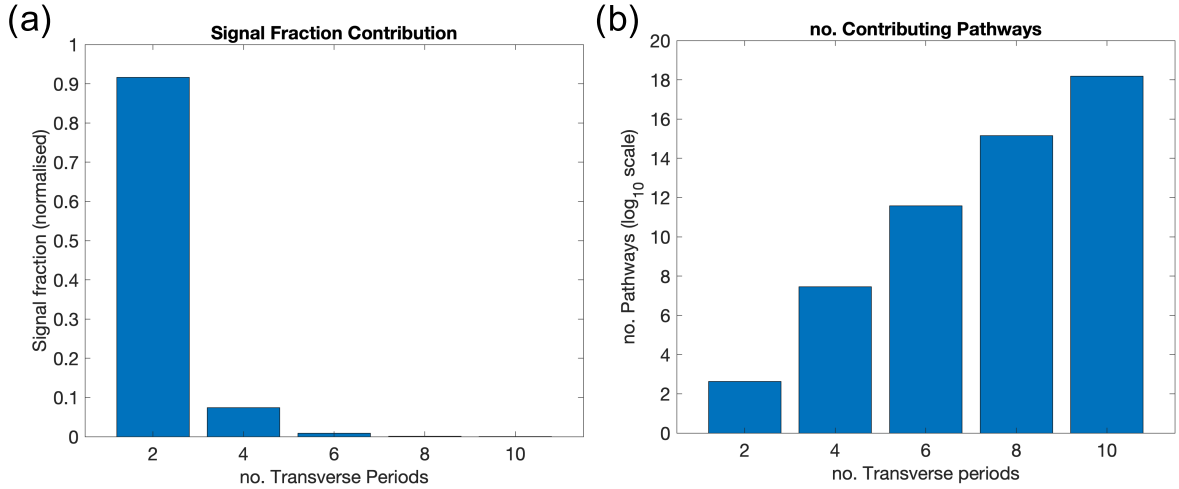


Figure S6: **DW-SSFP signal as a function of the number of transverse periods experienced by the magnetisation**. Based on the default modelling parameters and implementation thresholds defined in the Methods, we observe a decreased fraction of signal associated with pathways that experience a greater number of TRs in the transverse plane (a), associated with an increased number of pathways (b). Taken together, the default implementation of the time-*independent* pathway framework is associated with $>10^{18}$ signal forming pathways, with approximately 99% of the signal associated with pathways that persist for four transverse periods or less. To explore how changing parameters influences the contribution of pathways, find the associated code here.

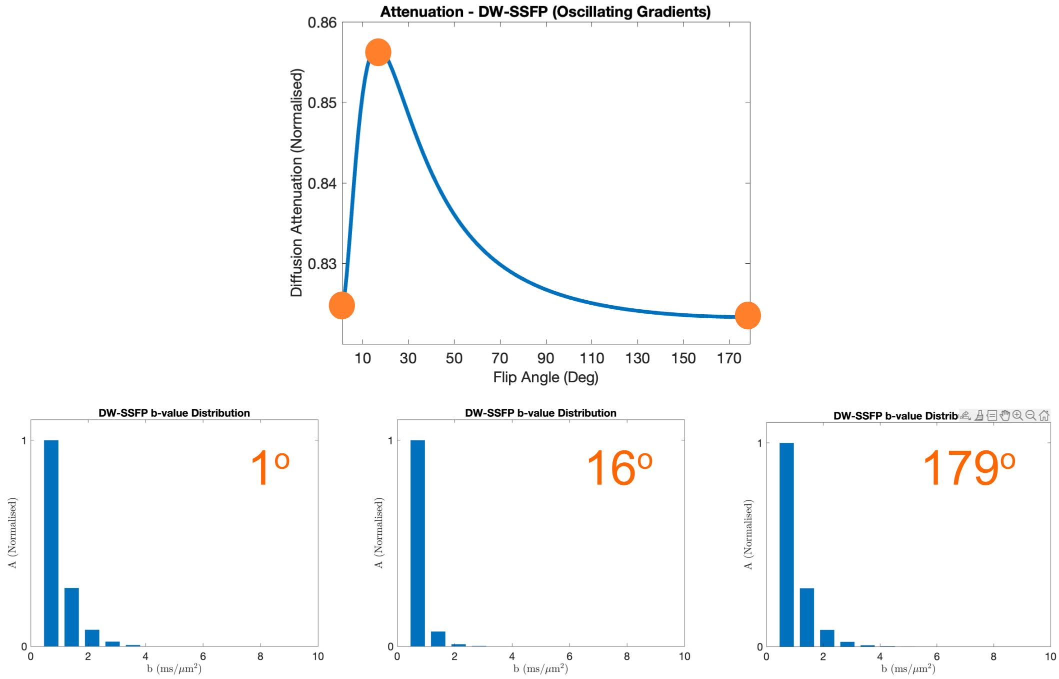


Figure S7: **Change in b-value weighting of the DW-SSFP signal with flip angle (oscillating gradients)**. Maximum diffusion attenuation with oscillating gradients is identified at both very low and high flip angles, in contrast to conventional DW-SSFP which sees increased attenuation at lower flip angles only. The b-value distributions at very low and high flip angles are almost identical, despite differences in the pathways that predominantly contribute to the signal. Specifically, this reflects the contribution of stimulated-echo pathways that persist for several periods in the transverse plane (very low flip angles) and spin-echo pathways that experience repeated pairs of dephasing and rephasing (very high flip angles). Parameters based on Aggarwal et al.[3] (see Methods). To explore how changing parameters influences the b-value plot, find the associated code here.

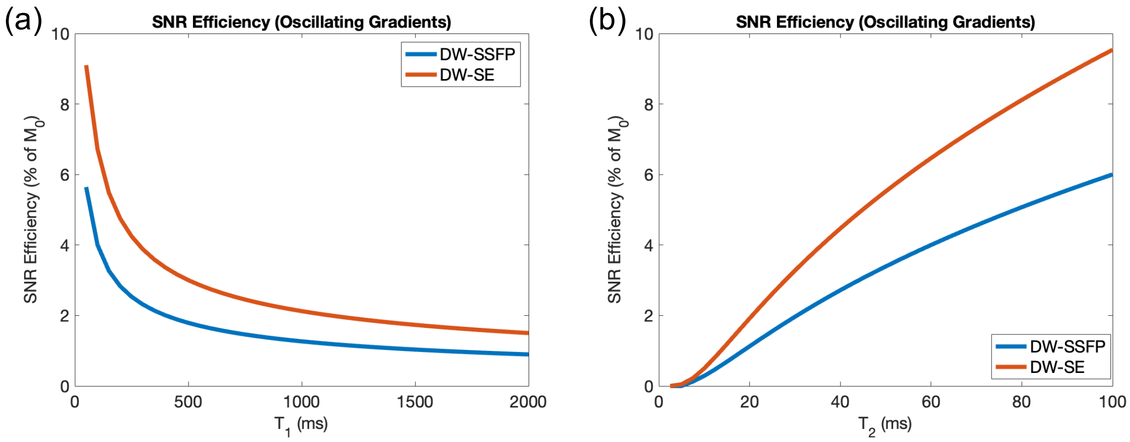


Figure S8: **SNR-efficiency properties of DW-SSFP (Oscillating Gradients) with relaxation**. SNR-efficiency estimation for DW-SSFP and DW-SE with oscillating gradients at a fixed encoding duration (10 ms per gradient) as a function of $T_1$ (a) and $T_2$ (b). DW-SE predicts higher SNR-efficiency versus the DW-SSFP sequence across all investigated regimes. Simulations based on mean estimates of $T_1$, $T_2$ & D from a cohort of post-mortem brains assuming free diffusion. For full information, see Methods. To explore how changing parameters influences these relationships, find the associated code here.

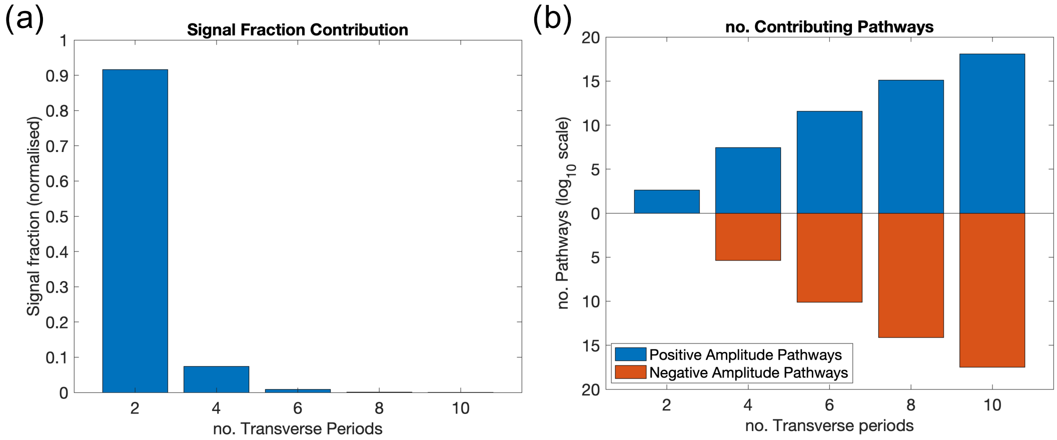


Figure S9: **DW-SSFP signal as a function of the number of transverse periods experienced by the magnetisation (positive and negative contributions)**. Based on the default modelling parameters and implementation thresholds defined in the Methods, we observe a decreased fraction of signal associated with pathways that experience a greater number of TRs in the transverse plane (a), associated with an (b) increased number of pathways. Whilst the contributed signal fraction is reduced as a function of the number of transverse periods (a), the relative signal fraction is formed by the cancellation of many pathways with positive (b – blue) and negative (b – red) amplitudes. For example, whilst the signal fraction associated with 10 transverse periods is ~0.02% of the total signal, it is formed from the weighted sum of ~$10^{18.1}$ positive amplitude pathways and ~ $10^{17.5}$ negative amplitude pathways.

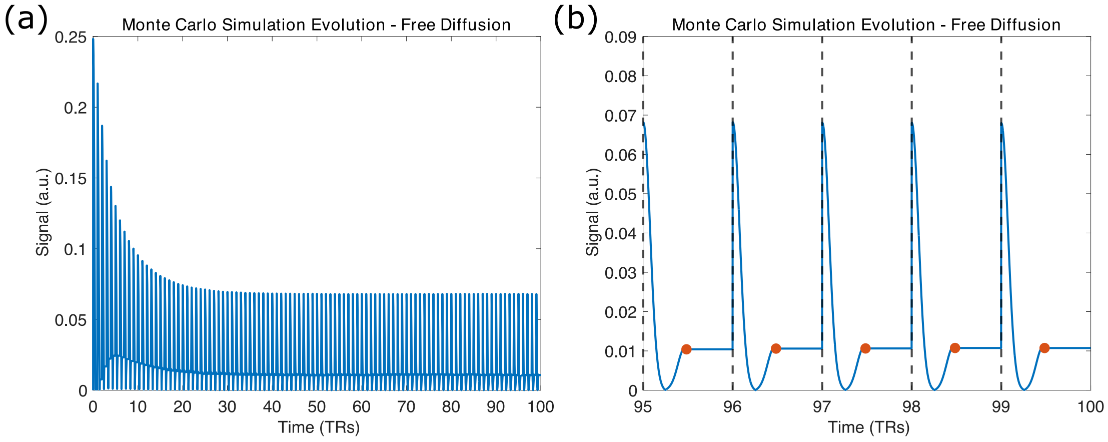


Figure S10: **Evolution of the DW-SSFP signal**. As is the nature of steady-state sequences, magnetisation evolves for many TRs until it reaches a steady state. (a) displays the evolution of the DW-SSFP signal for 100 TRs derived from a Monte Carlo simulation of free diffusion at 30° (see Methods). The simulation reaches a steady state after 10s of TRs. (b) displays a zoom of the final 5 TRs of the Monte Carlo simulation once a steady state has been reached. The dashed black lines represent the position of RF pulses, with the orange dots indicating the end of the applied diffusion gradient ($T_1$ and $T_2$ relaxation modelled instantaneously at the start of each TR).
